# Pif1-family helicases cooperate to suppress widespread replication fork arrest at tRNA genes

**DOI:** 10.1101/082008

**Authors:** Joseph S. Osmundson, Jayashree Kumar, Rani Yeung, Duncan J. Smith

## Abstract

*Saccharomyces cerevisiae* encodes two distinct Pif1-family helicases – Pif1 and Rrm3 – which have been reported to play distinct roles in numerous nuclear processes. Here, we systematically characterize the roles of Pif1 helicases in replisome progression and lagging-strand synthesis in *S. cerevisiae*. We demonstrate that either Pif1 or Rrm3 redundantly stimulate strand-displacement by DNA polymerase δ during lagging-strand synthesis. By analyzing replisome mobility in *pif1* and *rrm3* mutants, we show that Rrm3, with a partially redundant contribution from Pif1, suppresses widespread terminal arrest of the replisome at tRNA genes. Although both head-on and codirectional collisions induce replication fork arrest at tRNA genes, head-on collisions arrest a higher proportion of replisomes; consistent with this observation, we find that head-on collisions between tRNA transcription and replisome progression are under-represented in the *S. cerevisiae* genome. Further, we demonstrate that tRNA-mediated arrest is R-loop independent, and propose that replisome arrest and DNA damage are mechanistically separable.

## INTRODUCTION

The entire genome must be replicated during each S-phase to avoid genome instability and ensure accurate transmission of genetic and epigenetic information. Eukaryotes initiate DNA replication from multiple origins per chromosome^1^,^2^. Following origin activation, the replisome faces the formidable task of unwinding and replicating tens- to hundreds of kilobases of DNA, which may contain various impediments such as ongoing transcription, stable protein-DNA complexes, DNA lesions, and intramolecular DNA secondary structures^3^. Replication fork blockage at such impediments can lead to fork collapse and DNA breaks^4^; however, such arrest can be detrimental even if fork integrity is maintained. Most intuitively, the arrest of two convergent replication forks bordering a region without a licensed replication origin will preclude timely replication of this region, potentially leading to under-replication and attendant downstream problems such as chromosome mis-segregation^5^. Therefore, continued replication fork progression on challenging templates is fundamentally important for cellular and organismal viability, even if the replication fork is able to progress past lesions, which can be repaired after S-phase is complete^6^.

Pif1 helicases are a conserved family of 5’-3’ helicases capable of removing proteins from DNA, and of unwinding duplex DNA, RNA:DNA hybrids, and G-quadruplexes – kinetically and thermodynamically stable intramolecular DNA secondary structures resulting from Hoogsteen base pairing between four planar guanine bases^7^–^10^. While most metazoans encode only one Pif1, *S. cerevisiae* encodes two separate helicases: *PIF1* and *RRM3*, which are proposed to have largely distinct functions. Pif1 inhibits telomerase^11^ and helps to prevent genome rearrangements at G-quadruplexes^10^. In addition Pif1 has been identified as an accessory helicase for the lagging-strand polymerase, DNA polymerase δ (Pol δ), through both genetic^12^,^13^ and biochemical experiments^14^. In contrast, Rrm3 has predominantly been implicated as a replisome component^1^,^15^ that facilitates passage past protein-DNA complexes including tRNA genes and centromere^16^ although more recently it has been found to contribute to the maintenance of G-quadruplex-forming structures in combination with Pif1^10^.

We have previously described the sequencing of Okazaki fragments, enriched by repression of DNA ligase I, as a high-resolution method for the analysis of Okazaki fragment maturation, chromatin assembly, and replication fork movement throughout the genome^17^,^18^. Here, we report the systematic and genome-wide analysis of the effects of both Pif1 and Rrm3 activity on replisome progression and lagging-strand synthesis in *S. cerevisiae*. We determine that Pif1 and Rrm3 act redundantly to stimulate strand-displacement by DNA polymerase δ during Okazaki fragment maturation. While previous work found a role for Rrm3 but not Pif1 in aiding in fork progression at tRNA genes^16^, we demonstrate that Pif1 does help to facilitate replisome mobility in cells lacking Rrm3. The absence of both helicases leads to a high rate of terminal replication fork arrest (hereafter referred to as arrest and distinguished from transient replisome pausing, which we refer to as stalling) at tRNA genes (hereafter referred to as tDNAs). Furthermore, we show increased arrest in response to head-on collisions, and provide evidence that tDNAs in *S. cerevisiae* show a directional bias that reduces the likelihood of such interactions. In contrast to previous reports, we do not detect robust and significant arrest at other likely substrates of Pif1 helicases, e.g. G-quadruplexes and highly transcribed RNA polymerase II genes. Surprisingly, we find that conditions reported to increase or decrease the levels of R-loops at tDNAs^19^ do not impact replisome arrest at these loci, suggesting that these structures do not impede replisome mobility in the context of replication-transcription conflicts at tDNAs.

## RESULTS

### Assaying lagging-strand synthesis and replisome progression using Okazaki fragment sequencing

Rrm3 has been shown to aid in replication fork progression through various difficult-to-replicate sites; genetic interactions also imply a role for Pif1, but not Rrm3, in lagging strand biogenesis^12^. Because of their role in fork progression and lagging strand synthesis, we used Okazaki fragment sequencing as an unbiased, genome-wide assay both for fork progression and lagging strand maturation in strains lacking either or both Pif1-family helicase(s). Previous work using a repressible DNA ligase to enrich unligated Okazaki fragments *in vivo* showed remarkably uniform replication fork progression through the genome in wild type cells^17^,^18^, in agreement with ChIP-chip time course data^20^. The use of asynchronous cultures ensured full Okazaki fragment coverage and allowed us to analyze replication throughout the entire genome^17^. Neither *PIF1* nor *RRM3* is essential for viability in *S. cerevisiae*, but Pif1 is needed for stable maintenance of the mitochondrial DNA. To avoid potential artifacts due to mitochondrial defects, we used the well-characterized *pif1-m2* allele, which maintains the mitochondrial function without detectable Pif1 in the nucleus^21^–^23^. *pif1-m2 rrm3Δ* double mutant strains are viable but grow slowly and rapidly accumulate suppressors. Consequently, all experiments were carried out using freshly streaked cells that had undergone a minimal number of divisions prior to being stocked.

We combined mutants in the Pif1-family helicases (*rrm3Δ, pif1-m2, rrm3Δ pif1-m2)* with a previously described doxycycline-repressible *CDC9* (DNA ligase I) allele to sequence Okazaki fragments genome-wide^18^ (Figure 1A). Lagging-strand sequencing can be used to directly infer replication direction because Okazaki fragments are synthesized on the Watson strand by a leftward-moving fork and on the Crick stand by a rightward-moving fork^17^ (Figure 1A). Origin usage, detectable as an abrupt transition from leftward-to rightward-moving forks, appears similar between all four strains (Fig. 1A and Figure S1), and fork progression appears largely unaltered in the *rrm3Δ* and *pif1-m2* strains. However, the double mutant shows differences in Okazaki fragment distributions between replication origins, suggestive of significantly altered replication fork progression (Fig. 1A – gray boxes represent regions in which Okazaki fragment distributions are visibly different between wild-type and *rrm3Δ pif1-m2* mutant).

**Figure 1:**
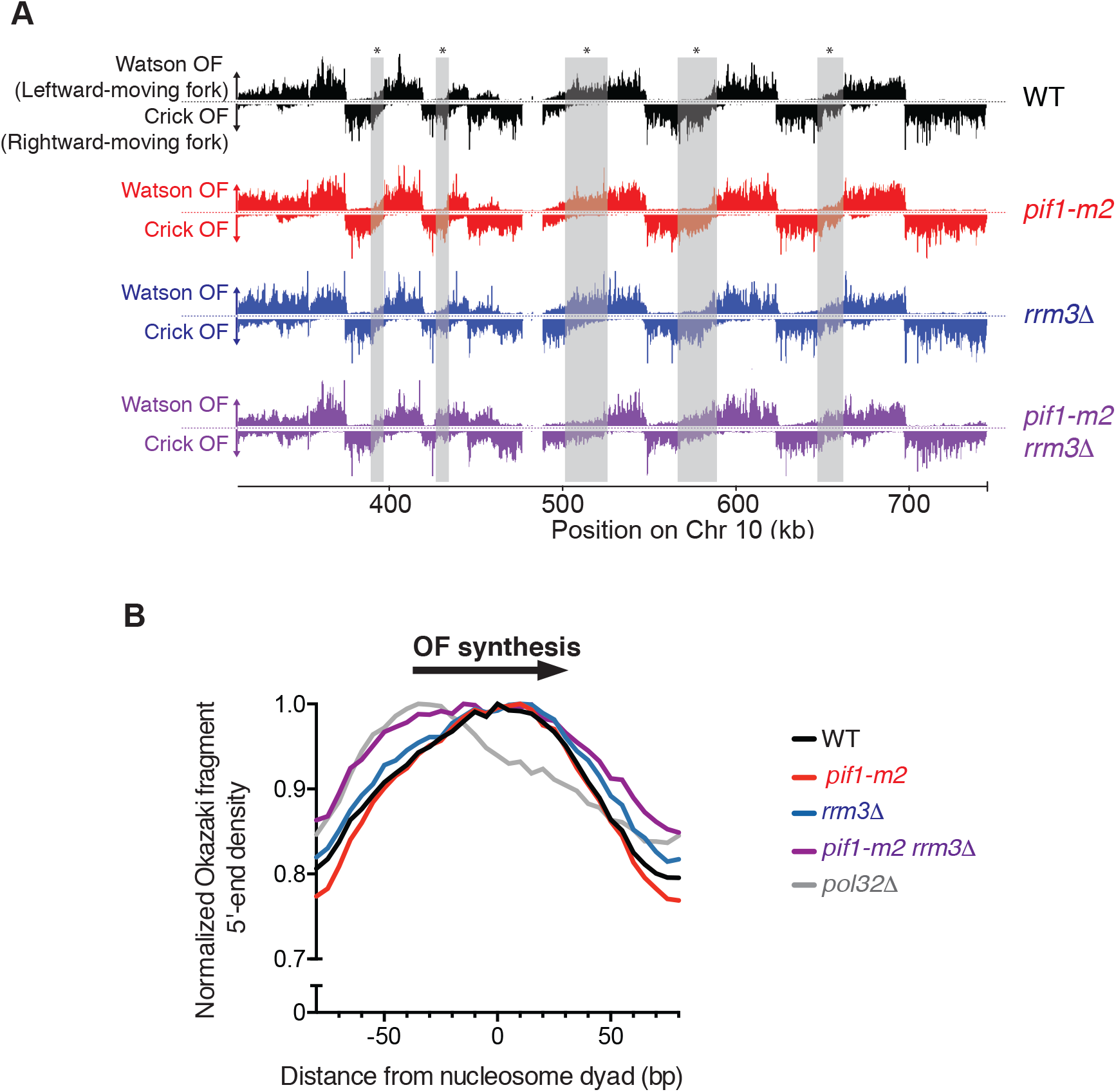
Okazaki fragment Sequencing is a quantitative and genome-wide assay for replisome mobility and lagging strand biogenesis in WT and mutant cells. (A) Distribution of Watson- and Crick-strand Okazaki fragments across the right arm of chromosome 10 in wild-type, *rrm3Δ*, *pif1-m2*, and *pif1-m2 rrm3Δ S. cerevisiae* strains. Watson strand fragments result from leftward-moving replication forks and are shown above each axis; Crick strand fragments result from rightward-moving forks and are shown below the axis. Grey boxes indicate regions where differences between distributions in wild-type and *pif1-m2 rrm3Δ* can be readily observed. Data were visualized using Mochiview (Homann and Johnson, 2010). (B) Either Pif1 or Rrm3 is required for normal DNA Pol δ displacement synthesis through nucleosomes. Distribution of Okazaki fragment 5’ termini from the indicated strains around consensus *S. cerevisiae* nucleosome dyads ^59^. Data are normalized to the maximum value in range and binned to 5bp. Data from the *pol32Δ* strain are from Smith and Whitehouse, 2012.

### Either Pif1 or Rrm3 can act as a lagging strand processivity factor

Lagging-strand synthesis in eukaryotes involves the generation of 5’ flap structures via strand-displacement synthesis by DNA polymerase δ (Pol δ), and their cleavage by structure-specific nucleases^24^. We have previously shown that Okazaki fragment maturation occurs in the context of nascent nucleosomes, and that Okazaki fragment ends are enriched around nucleosome midpoints due to the limited ability of Pol δ to penetrate the protein/DNA complex.^18^ Nucleosome penetrance during Okazaki fragment processing, which is reduced in strains lacking *POL32,*^18^ is a proxy for the level of Pol δ processivity during strand-displacement synthesis. In *pif1-m2* and *rrm3Δ* single mutants, the distribution of Okazaki fragment 5’ and 3’ termini around all nucleosomes^25^ was similar to wild-type (Fig. 1B & S2A). However, in the double mutant strain, both 5’ and 3’ termini showed a pronounced shift toward the replication fork-proximal edge of the nucleosome (Fig. 1B & S2A, purple line) similar to the effect of deleting *POL32.* These data are consistent with a reduction in the ability of Pol δ to carry out strand displacement on nucleosomal templates throughout the majority of the *S. cerevisiae* genome, and argue that either Pif1 or Rrm3 is required for processive Pol δ synthesis and wild type-like lagging strand synthesis. Okazaki fragments across all strains were similarly sized and equivalently poised for ligation (Fig. S2B), suggesting that the reduction in strand displacement seen in *pif1-m2 rrm3Δ* cells does not result in a substantial inability to complete lagging-strand processing. We therefore conclude that either Rrm3 or Pif1 is sufficient to act as a processivity factor for DNA Pol δ during lagging-strand biogenesis – i.e. the two helicases are completely redundant in this context (Fig. S3 and see discussion).

### tDNAs act as point replication terminators in the absence of Rrm3, and Pif1 can serve as a backup helicase in the absence of Rrm3 activity

A point replication origin can be identified from Okazaki fragment sequencing data as an abrupt reduction in Watson-strand coverage (moving from left to right) with a corresponding increase in Crick-strand coverage; the magnitude of this change is proportional to the efficiency of origin firing^17^,^26^. Analogously, a point replication terminator will manifest as the converse change, with Watson signal increasing at the expense of Crick signal (Figure 2A). Raw Okazaki fragment sequencing data from the *pif1-m2 rrm3Δ* double mutant contained numerous strand transitions suggestive of termination (Fig. 2B). We noticed that these apparent terminators often occurred close to tDNAs and, because these are known sites of fork pausing in the absence of Rrm3^1^,^16^, we undertook a systematic and quantitative analysis of fork progression, stalling, and arrest at all tDNAs genome-wide and in all four strains.

**Figure 2:**
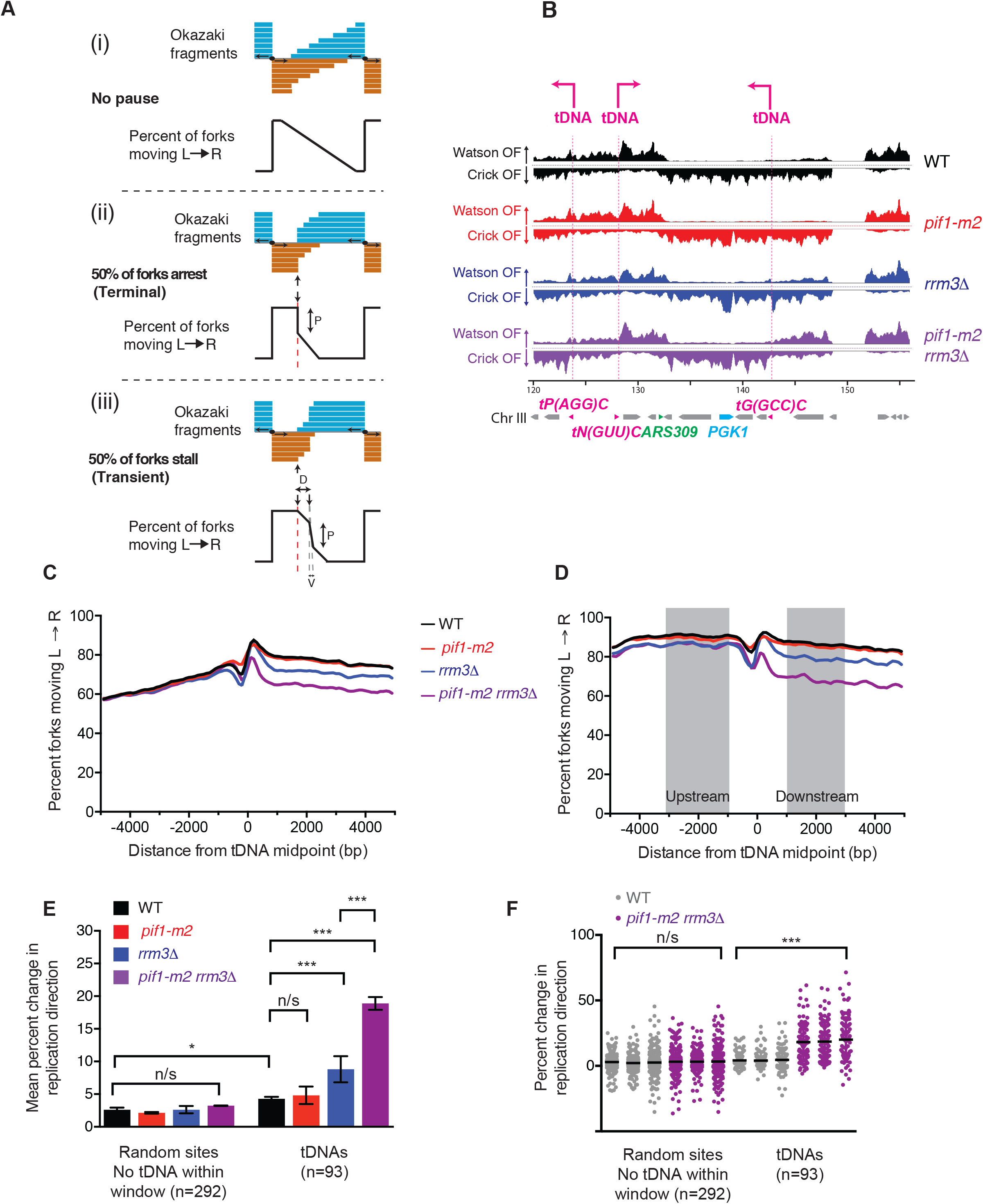
tDNAs are sites of replication fork arrest in *rrm3Δ* and *rrm3Δ pif1-m2* strains. (A) Schematic depicting the expected Okazaki fragment distributions (lower panels, orange and blue) and the percent forks moving left-to-right (upper panels, black line) through a region that (i) allows unimpaired replisome movement, (ii) terminally arrests the replisome, or (iii) transiently stalls the replisome. The magnitude of the strand transition, P, is proportional to the number of forks stalling or arresting: in the event of a transient pause, this transition will be offset from the element by a distance, D, and broadened over a window, V, dependent on pause duration and replication fork speed. (B) Distribution of Waston- and Crick-strand Okazaki fragments across an annotated 40 kb region of *S. cerevisiae* chromosome 3 as in Figure 1A. Locations of tRNA genes (tDNAs) are shown as pink lines and are indicated above the sequencing data. (C) Analysis of replisome direction around all 275 tDNAs in the *S. cerevisiae* nuclear genome. tDNAs replicated predominantly by rightward- or leftward-moving forks were analyzed separately, and the data superimposed such that the direction of fork motion is always from left to right (see methods). (D) Analysis of replisome direction, as in C, at the 93 tDNAs without a replication origin or sequence gap within ± 5 kb. The percent change in replication fork direction between the indicated regions 1-3 kb up- and downstream of a genomic meta-element is used to calculate termination signal in subsequent analyses (see methods). (E) Grand mean ± SD of the change in replication direction, interpreted as indicative of replisome arrest, at tDNAs and similarly filtered random sites in three independent biological replicates. Significance was determined by Monte Carlo resampling; ; *** indicates a p<0.0001; * indicates 0.0001 < p < 0.05; n/s indicates p>0.05. (F) One-dimensional scatter plot of the change in replication direction at the 93 tDNAs and 292 random sites in three biological replicate samples from E. Termination signal was calculated for each site as in part E and Black bars indicate the mean of each dataset. Significance was determined by Monte Carlo resampling; *** indicates a p<0.0001

The expected Okazaki fragment distribution around a a site of replication fork arrest (terminal) or stalling (transient) affecting 50% of replication forks is shown schematically in Fig. 2A. Only arrest or very long-lived replisome stalling events – those of sufficient duration to allow complete replication of the region downstream of the element by a convergent replisome – should give rise to distributions that show abrupt strand transitions directly at the termination element but are otherwise unaffected up-or downstream (compare Figure 2A(ii) and (iii)). By contrast, stalls should generate strand transitions that alter the observed sites of replisome convergence but are offset from the site of stalling. We note that our definition of replisome arrest does not necessarily imply that the replisome is unable to restart, but rather that such restart is not observed in the time taken for a convergent fork to complete replication.

To investigate replisome stalling or arrest at all 275 nuclear tDNAs on a single plot, we separately analyzed Okazaki fragment distributions around tDNAs predominantly replicated by leftward- or rightward-moving forks (see methods and Fig. S4A), and reversed the data from the leftward-moving set such that all replication forks were effectively moving from left to right as schematized in Fig. 2A. Meta-analysis of effective replication fork direction around all tRNA genes reveals a dramatic change in fork mobility in the *rrm3Δ* and *pif1-m2 rrm3Δ* strains (Fig. 2C), consistent with a high degree of replisome arrest at tDNAs. The transition was sharp and centered on the meta-tDNA midpoint – the predicted behavior for a site of fork arrest as opposed to a termination zone or stall (Figure 2C). However, the steady change in replisome direction observed upstream of the meta-tDNA, which is due to proximal replication origins (tDNAs are located, on average, near replication origins, Figs. S4 B-D and Table S1 and S2), precluded the direct quantitation of these changes in replisome mobility. To avoid this complication, we selected a stringent set of 93 tDNAs for quantitative analysis, excluding those with either an active replication origin^17^ or a sequence gap within the region ±5 kb from the tDNA midpoint (see Methods and Table S1). While we use this stringent set of tDNA sites for quantification, we note that there is clear evidence for fork arrest at essentially all 275 tDNAs in the *pif1-m2 rrm3Δ* strain (Figure 2C and S5A). Meta-analysis of Okazaki fragment distributions around this stringent tDNA set (Fig. 2D) allows us to calculate the mean change in replication direction, which serves as a direct and quantitative readout of the proportion of replisomes arresting at the meta-element genome-wide.

Because of fluctuations in sequence coverage in the immediate vicinity of tDNAs, we calculated the mean replication direction in the regions 1-3 kb up- and downstream of the tDNA midpoint (Fig. 2D), and subtracted the latter from the former to calculate the proportion of replisomes terminating within ±1 kb of the each tDNA in each dataset. We calculated the grand mean from datasets obtained from three biological replicates (Figure 2E) and evaluated significance by Monte Carlo resampling. The WT and *pif1-m2* strain show no significant difference, whereas the *rrm3Δ* strain shows a significant (p<0.0001), arrest and the *pif1-m2 rrm3Δ* double mutant shows robust and significant (p<0.0001) arrest with nearly 20% of replication forks arresting and being rescued by a fork arriving in the opposite direction (Figure 2E). Mean values across the three datasets were highly reproducible (Fig. 2F). Therefore, we conclude that Rrm3 normally suppresses replisome arrest at tDNAs. Surprisingly, Pif1 can partially compensate for the absence of Rrm3 at tDNAs, and in the absence of both helicases, replication termination at tDNAs is extremely pronounced.

To determine whether the observed replisome arrest was specific to tDNAs, we analyzed replication direction around a set of random sites filtered equivalently to our tDNA set to remove those with proximal replication origins and/or sequence gaps (Table S1). In addition, due to the strong stalling signal at tDNAs, we excluded sites within 5 kb of a tDNA midpoint (see below and Figure S4H). Quantitation of replisome stalling revealed no replication termination above WT level in any mutant strain at random sites (Fig. 2E-F & S4E-H). The overall lower unidirectionality of fork movement at random sites and tDNAs in the double mutant strain is due to forks that have previously encountered a replication origin proximal tDNA, as the difference is no longer evident if we remove sites with an upstream tDNA and correct for the slight (5%) difference in origin efficiency (compare Figure S4C-D, F-G). There is a small (≈2%) but significant (p=0.02, Fig. 2E) increase in fork arrest at tDNAs compared to random sites in WT cells, in agreement with previous estimates that ~1% of replisomes terminally stall at tDNAs in wild-type yeast^20^. Subsequent analysis of tDNA orientation dependence (Fig. 5) is consistent with these findings.

### Pif1 helicases prevent replication fork arrest at essentially all tDNAs

We next sought to examine each tDNA separately, and to again analyze all 275 tRNA genes, to determine whether all tDNAs are capable of provoking arrest of the replication machinery in the absence of both Pif1-family helicases. At each tDNA, the difference in Okazaki fragment strand bias between WT and the *pif1-m2 rrm3Δ* strain is minimal before the fork reaches the tDNA. In leftward moving forks (Figure S5A: bottom of heatmap), there is a substantial decrease of the proportion of forks moving leftward after the tDNA midpoint. The inverse is true at essentially all tDNAs replicated by a rightward moving fork (top of the heatmap); to the left of the site, before the fork has encountered the tDNA, there is little difference between the WT and the *pif1-m2 rrm3Δ* mutant strain, but after the fork has encountered the tDNA, the proportion of rightward moving forks is lower due to arrested forks that are rescued by a leftward moving fork (Figure S5A). Moreover, comparing the termination signal at all 93 tDNAs used in our previous analysis in the double mutant to the wild-type strain indicates substantial arrest at almost every site, reproducible across multiple datasets (Figure S5B).

### Validation of replication fork arrest at tDNAs by 2-D gel

To validate the use of Okazaki fragment sequencing to map replication fork arrest, we directly assayed replication fork mobility around a single tDNA by neutral-neutral 2-D gel agarose electrophoresis. We identified a site on chromosome III where a rightward moving replication fork stalls at a tDNA oriented head-on with respect to replication (Figure 2A). Previous work has identified the adjacent highly transcribed RNA Pol II gene^27^ as a replisome stalling element in *rrm3Δ* cells through increased occupancy of DNA Polymerase ε. We cloned the origin that drives replication toward this site (ARS309) and the tDNA itself (*tG(GCC)C*) into a plasmid such that replication and transcription should occur in the head-on orientation as at the native locus (Fig. 3B). We isolated plasmids from asynchronous cultures of all four strains, digested with EcoRI to isolate the left half of the plasmid, and probed the AMP^R^ gene, which is unique to the plasmid (Fig. 3C). Stalling or arrest of the leftward-moving replication fork are predicted to give rise to a dark spot on the Y-arc; arrest or long-lived stalling of the leftward-moving fork should allow the rightward-moving fork to progress past the second EcoRI site, generating double-Y structures in the context of stalling or a specific X spot in the context of fork arrest (Fig. 3A).

**Figure 3:**
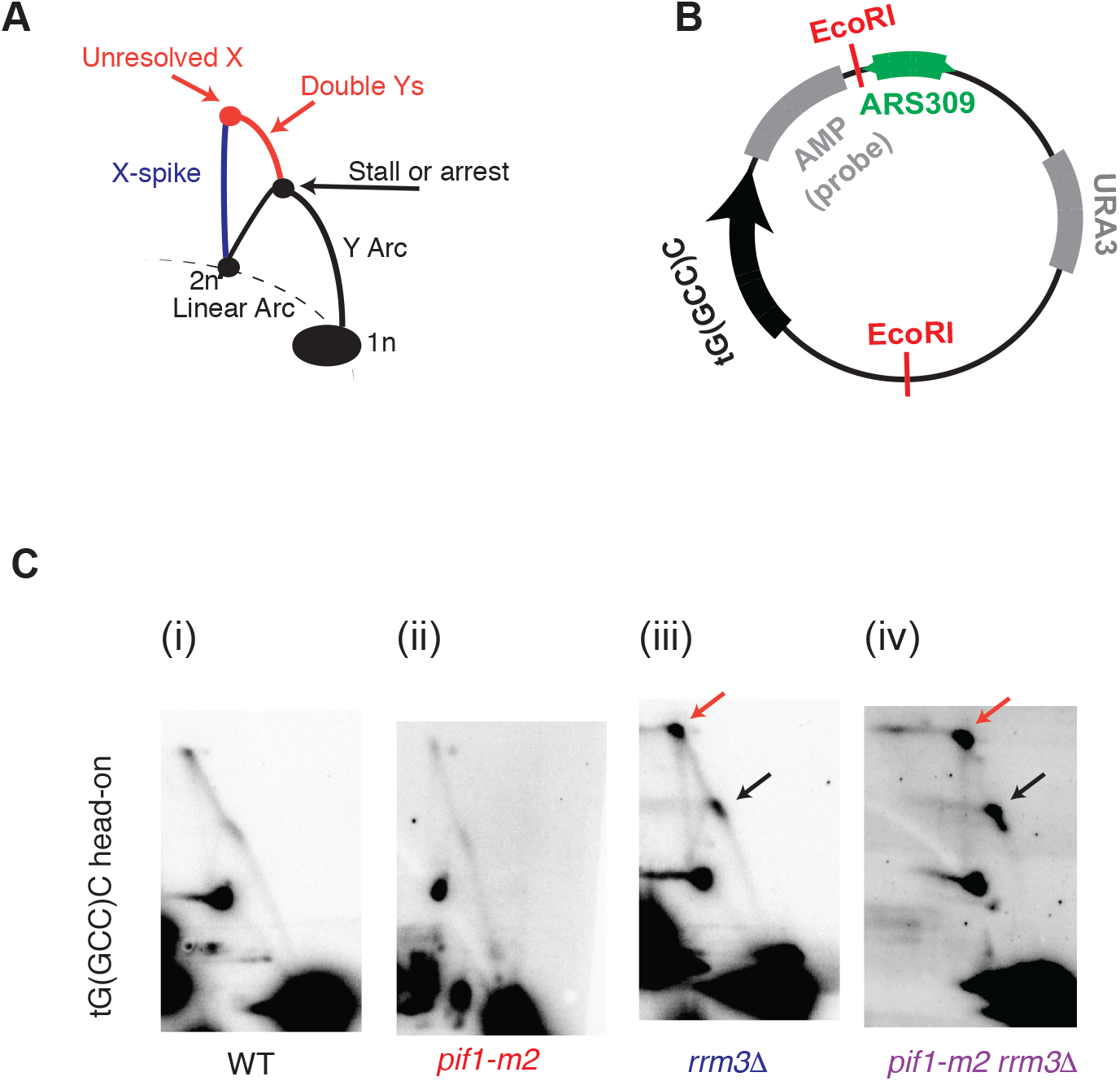
2-dimensional gel validation of replication fork arrest in *rrm3Δ* and *rrm3Δ pif1-m2* strains. (A) Schematic of intermediates, resolved in a neutral-neutral 2-dimensional agarose gel, resulting from the replication of plasmid (B) in asynchronous cultures. (C) Neutral-neutral 2-Dimensional gel analysis of the indicated plasmid in WT, *pif1-m2*, *rrm3Δ* and *pif1-m2 rrm3*Δ cells, as indicated. 2-D gel signals consistent with replisome stalling and/or arrest are indicated in colors corresponding to the schematic in Fig. 3A. Plasmids were cut with the indicated restriction enzymes and crosslinked with psoralen prior to electrophoresis.

In agreement with our genome-wide Okazaki fragment sequencing data, we see a slight or no enrichment of replication intermedites on the Y-arc in the WT and *pif1-m2* cells, a moderate enrichment in the *rrm3Δ* cells, and a strong enrichment in the *pif1m2-rrm3Δ* cells (Figure 3C, black arrows) suggesting that Pif1 can indeed act as a backup for Rrm3 activity. In all cases, and consistent with arrest rather than stalling, an X-spot signal is observed (Figure 3C, red arrows). Thus, 2D gel analysis further supports our conclusion that Rrm3 suppresses replisome arrest at tDNAs, and that Pif1 can partially compensate for its absence.

### Highly transcribed RNA Pol II genes and G-quadruplexes do not induce significant replisome arrest in *pif1* and/or *rrm3* mutant strains

Previous work has implicated Pif1, Rrm3, or both proteins in aiding replication fork progression through highly transcribed RNA Polymerase II (RNAP2) genes^1^,^16^,^27^ and G-quadruplex structures *in vivo*^10^,^28^. Therefore, we analyzed Okazaki fragment distributions to assay replication stalling or arrest around highly transcribed RNAP2 genes (from ^29^, Table S4), and the ribosomal protein genes as a representative group of coordinately regulated, highly transcribed RNAP2 genes (from ^30^ Table S4), and G-quadruplexes (from ^31^, Table S5), selected in the same way as our 93 stringent tDNAs (i.e. no origin of replication or sequence gap within 10kb of the site).

When we did not remove sites that overlap with a tDNA, a small but significant arrest was detected in highly transcribed RNA Pol II genes (p= 0.01) and G-quadruplex motifs (p=0.01). However, we also observed a small but significant signal at our randomly selected sites (p=0.006; Figure S6A). We therefore eliminated the small number (≈10%) of sites that overlapped with tDNAs, and the significant increase in arrest at all these classes of sites was no longer observed (random sites: p=0.11; RNAP2 genes: p=0.08; G4: p=0.06; compare Figure 4A and S6A).

**Figure 4:**
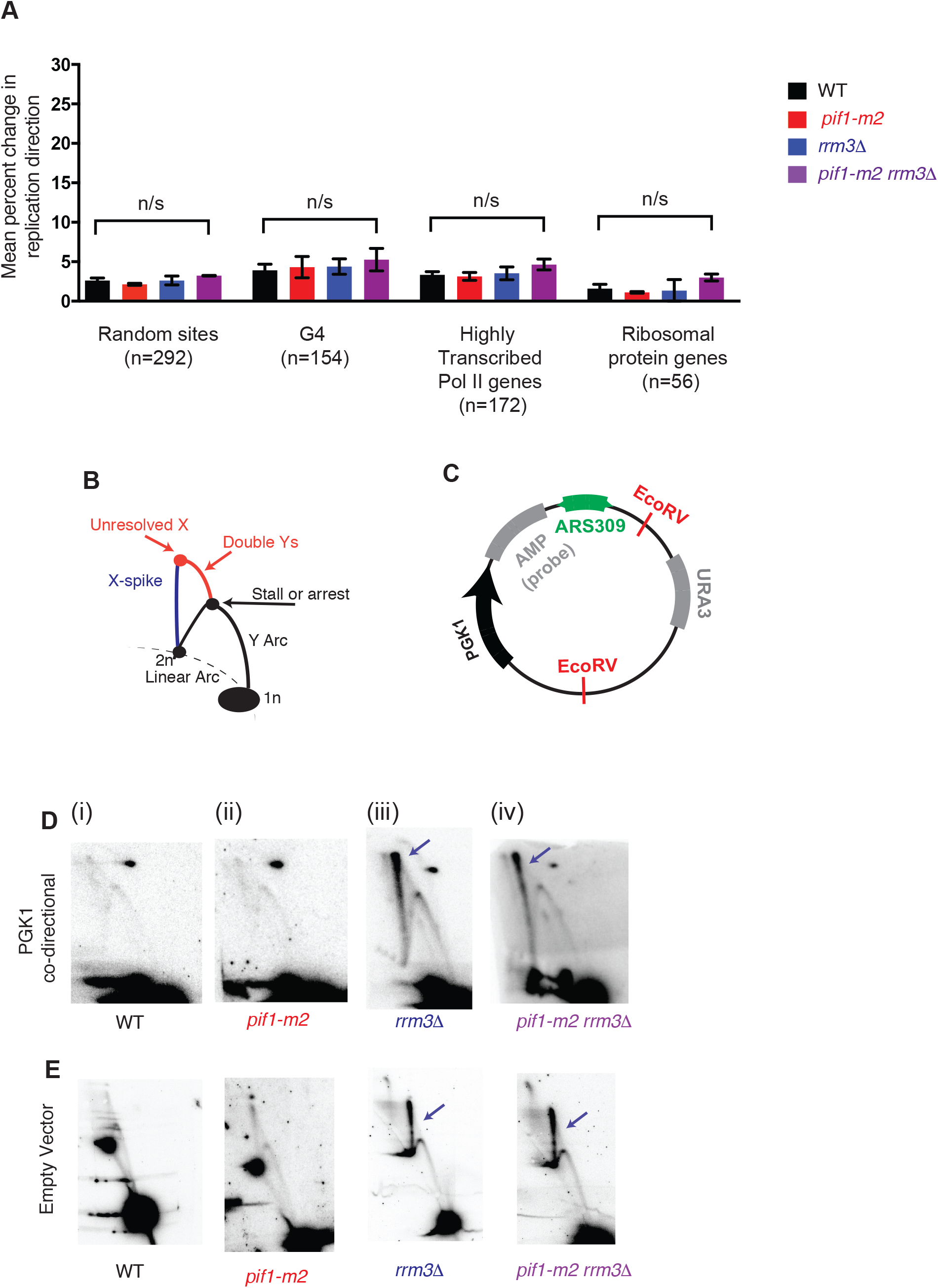
G-quadruplexes and highly transcribed RNA Polymerase II genes do not contribute to significant replication fork stalling or arrest genome wide. (A) Grand mean ± SD of the change in replication direction, interpreted as replisome stalling or arrest, around the indicated classes of genomic locus from three biological replicates as in Fig. 2E. Loci were filtered as for the tDNA analysis in Fig. 2D-F, excluding sites within ±5 kb of a sequence gap, replication origin, or tDNA. Significance was determined by Monte Carlo resampling; *** indicates a p<0.0001; n/s indicates p>0.05. (B) Schematic of intermediates, resolved in a neutral-neutral 2-dimensional agarose gel, resulting from the replication of plasmid (C) in asynchronous cultures. (D-E) Neutral-neutral 2-Dimensional gel analysis of a plasmid with or without a *PGK1* insert in asynchronous WT, *pif1-m2*, *rrm3Δ* or *pif1-m2 rrm3*Δ, as indicated. 2D gel signals consistent with changes in replication fork mobility or resolution are indicated in colors corresponding to the schematic in Fig. 3A. Plasmids were cut with the indicated restriction enzymes and crosslinked with psoralen prior to electrophoresis.

We considered many subsets of both RNA Pol II genes and G-quadruplex structures, including G-quadruplexes that were previously shown to be enriched in Pif1 binding^28^, and did not identify a biologically relevant subset that showed robust and significant stalling or arrest when sites that overlap with tDNAs were excluded (Figure S6C). Importantly, we find that tDNAs are not more likely than random sites to contain G-quadruplex forming sequences (Table S2). *pif1-m2* cells were previously shown by 2-D gel to have detectable replisome stalling only in the presence of hydroxyurea (HU), which depletes dNTP pools and thus slows or stalls forks depending on the concentration.^28^ However, even during growth in 25 mM HU, we see no evidence for significant terminal stalling at G-quadruplex sites genome wide (Figure S6D).

We did find a small (3.5%) but statistically significant (p = 0.002) arrest signal in the *pif1-m2 rrm3Δ* mutant compared to WT cells at G-quadruplex sites that were previously found to stall the fork by ChIP-chip of DNA Pol ε^28^. Previous work on fork progression through G-quadruplex motifs in the absence of Pif1 identified a small subset of G-quadruplexes that were enriched for replisome components, and another small subset of G-quadruplexes that showed evidence for Pif1 binding by ChIP-chip^28^. We asked if we could similarly extract a subset of sites that explained the fork arrest in some but not all G-quadruplexes in the *pif1-m2 rrm3Δ* double mutant by Okazaki fragment sequencing. Indeed, by comparing the three replicate datasets and identifying G-quadruplex motifs where the difference in arrest between the WT and *pif1-m2 rrm3Δ* datasets was greater than the standard deviation between the datasets, we identified 46 sites (without a tDNA) where the mean termination signal was significantly higher (p<0.0001) in the double mutant than WT cells (Fig. S6E). However, if we performed an identical analysis starting with the same number of random sites and extracted those that showed arrest in the double mutant in the same way (again excluding those with a tDNA), we identified 43 random sites from which we could infer statistical significance (p<0.0001) for arrest in the double mutant compared to the WT (Figure S6E). Importantly, neither the random sites nor the G-quadruplex sequences in these ‘arrest’ subsets were enriched for any genomic feature, including other sites predicted to stall the replication fork (Table S2). Therefore, in our assay, sequences predicted to form G-quadruplex structures act no differently than sequences chosen at random, and statistically significant signal can be extracted from the noise in, or the shape of, the deep sequencing data.

To validate the absence of replisome stalling or arrest at highly transcribed RNA Pol II genes, we cloned *PGK1* and its promoter into the previously described ARS309-containing plasmid for 2-D gel analysis from asynchronous cultures. *PGK1* was previously implicated as a site of replisome stalling by ChIP-chip of Pol ε^27^ but, unlike the adjacent *tG(GCC)C*, did not perturb Okazaki fragment distributions (Fig. 2B). We did not observe substantial effects on replisome mobility at *PGK1* by 2-D gel in any of the four strains analyzed (Figure 4D and cf. Fig. 3C – note that the spot on the upper right in all four blots is due to background hybridization of the probe). However, a strong X-spike signal was observed in both *rrm3Δ* and *pif1-m2 rrm3Δ* (Figure 4D, blue arrows). To test the hypothesis that this X-spike represented a defect in the resolution of convergent replication forks in these strains, and that the defect in replication termination was not due to conflicts with transcription, we transformed cells with a plasmid lacking *PGK1*, and whose left half is therefore transcriptionally silent; the X-spike is still apparent in both *rrm3Δ* and *pif1-m2 rrm3Δ* strains even when the replication fork encounters no transcribing RNA polymerase (Fig. 4E, iii & iv).

Any fork stalling or arrest at RNA Pol II genes or G-quadruplex structures is likely to show an orientation effect (e.g. leading-strand G-quadruplex structures have been shown to induce more genome instability than lagging strand sites^32^). However, we see no significant orientation effect for either RNA Pol II genes or G-quadruplex structures (Figure S7). Therefore, we conclude that our combined analysis of Okazaki fragment distributions and plasmid replication intermediates that replisome stalling or arrest at G-quadruplexes and RNAP2 genes is extremely rare even during replication stress in the absence of both Pif1 and Rrm3 function, and that Rrm3, but not Pif1, plays a role in the resolution of converged replisomes.

### Replication fork arrest at tDNAs is partially orientation dependent

Conflicts between the transcription and replication machinery replication fork have been described *in vitro* and *in vivo* and in bacterial and eukaryotic systems^33^–^37^. There have been conflicting reports about whether tDNAs impede the replication machinery in WT cells. tRNA genes have been found, by 2-Dimensional gel to act as replication fork blocks regardless of orientation the absence of Rrm3 ^1^,^16^, whereas the original report of tDNA-mediated stalling by Deshpande and Newlon found that only head-on tDNAs stalled or arrested the replisome in WT cells^38^. Our high-throughput analysis offers the opportunity to directly compare fork arrest at all tRNA genes in the same sample and assay.

To determine the extent to which fork arrest at tDNAs is orientation-dependent, we separated the 93 tDNAs in our analysis into those replicated head-on or co-directionally with respect to RNA Pol III transcription (Figure 5A and see methods). In all strains, head-on collisions were significantly more likely to arrest the replication fork than co-oriented collisions (p<0.0001 for all strains; Figure 4B), with head-on tDNAs in the *pif1-m2 rrm3Δ* double mutant being the subset of sites with the most widespread arrest. Interestingly, while we find that while the proportion of replisomes that arrest at tDNAs increases upon deletion of *RRM3*, and further on additional mutation of *PIF1*, the relative ratio of head-on vs co-directional arrest remains constant. For validation, we switched the orientation of the tDNA in the plasmid-based 2-D gel assay in the *rrm3Δ* strain; more arrest, reflected by more intense localized signal on the Y-arc and X-spot, is observed when the tDNA is in the head-on orientation (Figure 4C). We conclude that tDNAs in either orientation can lead to replisome arrest, but that this arrest is approximately twice as frequent at tDNAs oriented head-on relative to replication. While Pif1 helicases reduce the overall frequency of fork arrest, they do not preferentially impact one orientation relative to the other.

**Figure 5:**
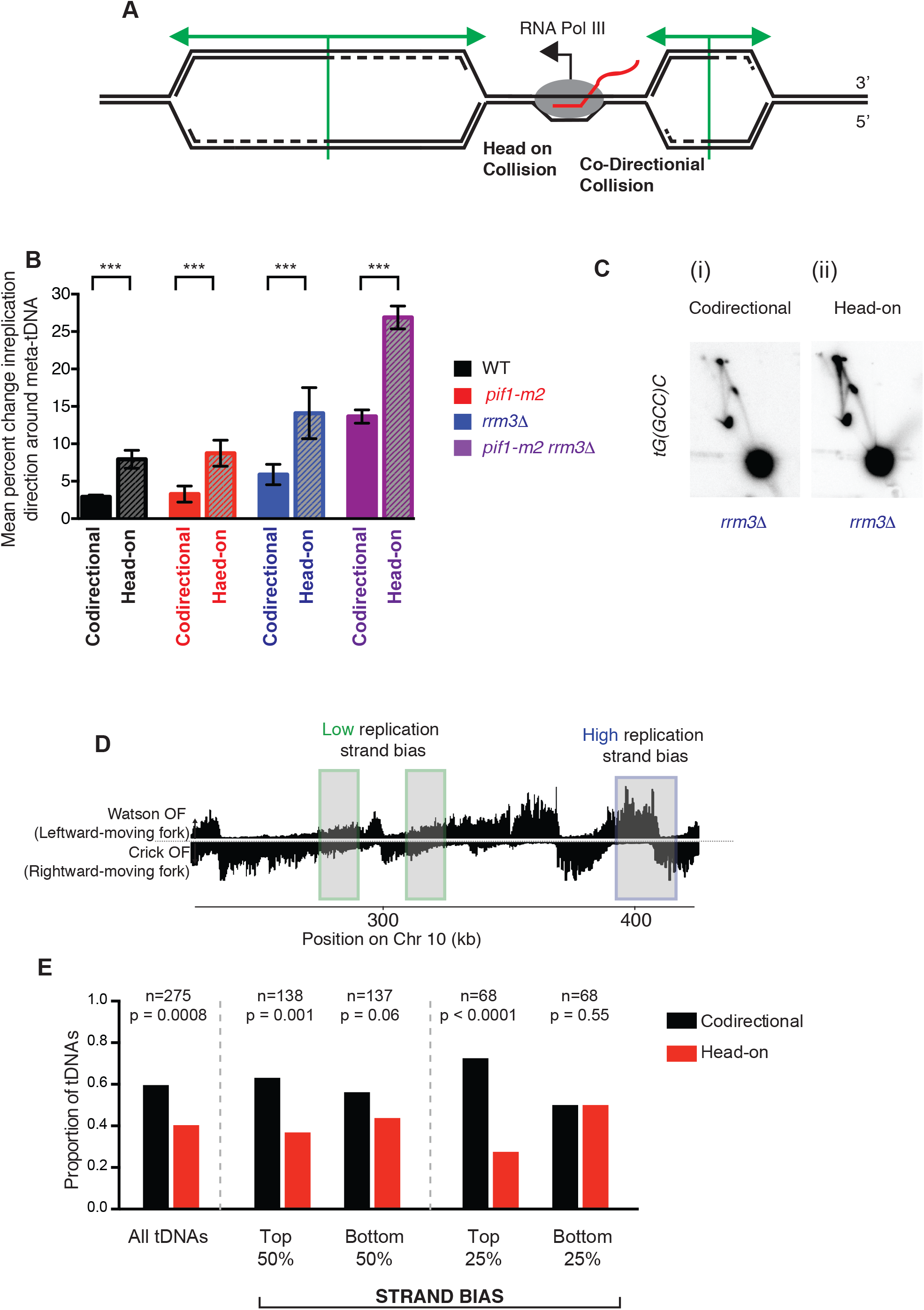
All tDNAs act as replication terminators, but head-on orientation between replication and transcription machinery increases fork arrest. (A) Schematic of two replication forks approaching a highly transcribed tDNA. The fork moving from left to right will meet the transcribing RNA Polymerase III molecule in a head-on fashion. (B) Grand mean ± SD from three biological replicates of the change in replication direction, interpreted as indicative of replisome arrest, at tDNAs transcribed co-directionally or head-on relative to the predominant direction in which they are replicated (see methods). Termination signal was calculated as in Figure 2E. Significance was determined by Monte Carlo resampling; *** indicates a p<0.0001 (C) Neutral-neutral 2-D agarose gel electrophoresis of the plasmid shown in Fig. 3B, but with *tG(GCC)C* in either the codirectional or head-on orientation as indicated, purified from an asynchronous *S. cerevisiae* culture. (D) Schematic of chromosomal regions of high (blue) and low (green) replication strand bias. (E) Bar plot indicating the proportion of tDNAs oriented head-on or codirectionally relative to replication, separated into bins (above or below the median strand bias for the 275 nuclear tDNAs) as indicated by strand bias as shown in D; p-values for cumulative binomial probabilities are indicated.

### Head-on collisions between tRNA transcription and replication are under-represented in the *S. cerevisiae* genome

To search for an evolutionary signature of increased replisome arrest due to head-on tDNA transcription, we interrogated tDNA orientation in regions of high replication strand asymmetry (Figure 5D). We examined all 275 tDNAs and find a significant overrepresentation of co-oriented transcription/replication conflicts (Figure 5E). We then sought to analyze tDNAs with high and low replication strand bias (see Figure 5D) to determine whether the likelihood of the conflict being always co-directional or head-on could predict the orientation of the tDNA. Indeed, in regions of high replication strand bias, where a tDNA is likely to be replicated in only one direction in almost all cells, there is an increased bias for co-oriented transcription/replication conflicts (Figure 5E). To show that the strand bias is predictive of tDNA orientation, we performed a logistic regression; increased strand bias leads to a significant decrease in the likelihood of a head-on orientation between transcription and replication (p=0.01, Odds ratio = −0.023). This observation, which to our knowledge represents the first demonstration of tDNA orientation bias in a nuclear genome, suggests an evolutionary pressure to co-orient tDNAs with replisome movement in *S. cerevisiae* and is consistent with the observation that head-on tDNAs are sites of significant replisome arrest even in wild-type cells. Equivalent analysis of protein-coding genes did not indicate a directional bias relative to replication, consistent with previous reports^39^ and our failure to detect directional fork stalling or arrest at these sites (Figure S7).

### R-loops do not significantly contribute to replication fork arrest at tDNAs

R-loops are RNA:DNA hybrid structures that form *in vivo* when the RNA transcribed by RNA Polymerase reanneals with the template strand, leaving the non-template strand of DNA extruded^19^,^40^. Importantly, R-loops are a directional feature for the replication fork: a Watson stranded gene would create an R-loop on the lagging strand for a leftward moving fork, and a Crick stranded gene would cause a leading-strand R-loop (see Figure 5A).

The RNA strand of R-loops is degraded by RNase H1; modulating the levels of RNase H1 and H2 in the cell can increase or decrease the level of R-loops^41^,^42^. In particular, a *rnh1Δ rnh201Δ strain* was recently shown by DRIP-seq using an anti-RNA:DNA hybrid antibody to have increased R-loop occupancy at tDNAs^19^. We sorted tDNAs by their previously reported levels of R-loops in wild-type cells (Figure 6A), binned tDNAs into quartiles, and analyzed replisome arrest by comparing Okazaki fragment distributions at high-DRIP (top quartile) vs low-DRIP (bottom quartile) tDNAs. Both tDNA subsets show strong replisome arrest in both *rrm3Δ* and *pif1-m2 rrm3Δ* strains (Fig. 6C). Direct quantitation of changes in replisome mobility at high- vs low-DRIP tDNAs is precluded by the distinct fork direction gradients observed upstream of the two subsets, presumably because the former lie, on average, closer to efficient replication origins. Therefore, we sought to modulate R-loop levels in the context of wild-type and *rrm3Δ* strains and directly assay the impact of this perturbation on replisome mobility. We constructed *rnh1Δ rnh201Δ* and *rnh1Δ rnh201Δ rrm3Δ* strains with repressible *CDC9*, and carried out triplicate replicates as in previous analyses. Additionally, we placed a highly inducible (at least 100-fold over endogenous, data not shown) *GAL1,10* promoter upstream of *RNH1* in *rrm3Δ* cells. Analysis of replication fork arrest in strains with increased or decreased RNaseH activity indicated no significant difference in either a wild-type or *rrm3Δ* background (Fig. 6D). Because increasing or decreasing the levels of R-loops at tDNAs does not exacerbate or alleviate fork arrest, we conclude that R-loops are extremely unlikely to underlie the orientation-based asymmetry of replisome arrest at tDNAs.

**Figure 6:**
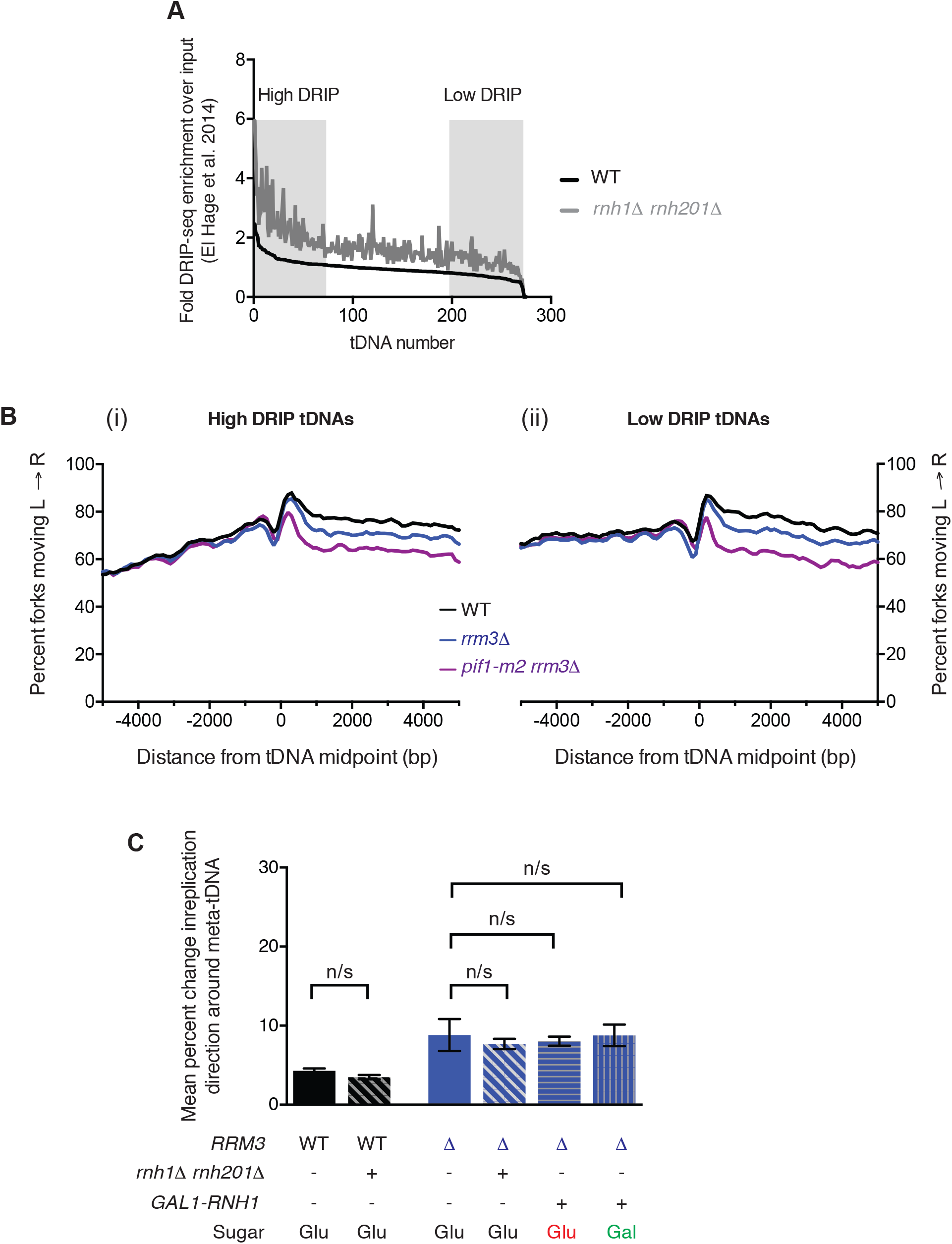
R-loops do not mediate replication fork arrest at tDNAs. A. R-loop load at all 275 tDNAs determined by DRIP-seq (El Hage *et al.,* 2014) in WT and *rnh1Δ rnh201Δ* cells. The level of R-loops over background was determined for a 400bp window around the tDNA midpoint. tDNAs were sorted by the level of R-loops in the WT strain and the levels in both the WT (black) and the *rnh1Δ rnh201Δ* double mutant (gray) were plotted. Regions corresponding to high- and low-DRIP in Fig. 6B are indicated. (B) Analysis of replisome direction around (i) high- and (ii) low-DRIP-seq tDNAs as identified in A. (C). Grand mean ± SD from three biological replicates of the change in replication direction, interpreted as indicative of replisome arrest, at the 93 tDNAs from Fig. 2E in the indicated strains. Cells were grown in either YEP + dextrose (Glu) or galactose (Gal) as specified. Significance was determined by Monte Carlo resampling; n/s indicates p>0.05.

## DISCUSSION

### Pif1 helicases and Okazaki fragment maturation

We show that Rrm3 is able to stimulate DNA polymerase δ strand displacement on the lagging strand (Figure 1). Previous genetic experiments implicated only Pif1 but not Rrm3 in lagging strand biogenesis: *pif1Δ* or *pif1-m2* act as suppressors of *dna2* mutations, and can restore viability to *dna2Δ* strains^13^ and Pif1 stimulates strand displacement by Pol δ *in vitro*^14^. These results have been taken as support for a two-nuclease model of Okazaki fragment processing in which Dna2 is responsible for the cleavage of Fen1-resistant long 5’-flaps generated by excessive strand displacement^24^. Conversely, *rrm3Δ* is synthetic lethal with *dna2* mutants^12^. Because Rrm3 is able to promote Pol δ processivity, we think it unlikely that genome-wide Okazaki fragment processing underlies the genetic interactions between *DNA2* and *PIF1/RRM3*. It is possible that the genetic interactions are due to telomere maintenance, as Dna2 and Pif1 both play critical roles in this process^11^,^43^. Additionally while Rrm3 can substitute for Pif1 during bulk Okazaki fragment processing, it cannot analogously replace Pif1 during Break-induced replication – another Pol δ-dependent synthesis reaction^44^; determining the molecular basis for this differential helicase requirement will provide important insights into both canonical and break-induced replication.

### tDNAs are the predominant site of replication fork arrest in cells lacking Rrm3 or Rrm3 and Pif1

As previously reported, we find that Rrm3 contributes significantly to replisome passage through tDNAs in *S. cerevisiae*, but we report a novel role for Pif1, which can partially compensate for the absence of Rrm3 at these sites. In the absence of both helicases, all tDNAs have the potential to stably or permanently arrest the replication fork. Indeed, the replisome arrest signal observed at tDNAs is sufficiently pronounced that analysis of replisome mobility at non-tDNA elements requires the removal of tDNA-proximal sites (see for example Figure S6H). Furthermore, the widespread fork arrest at tDNAs is apparent from the overall difference in fork unidirectionality in *pif1-m2 rrm3Δ* at random sites when the fork has already encountered an origin-proximal tDNA (Figure S4F versus G). While it is difficult to conclusively show that there is no tendency towards replisome arrest at any other site genome-wide in cells lacking Pif1 and/or Rrm3, the signal at tDNAs clearly reflects the vast majority of fork stalling or arrest genome-wide and is evident even in WT cells when the tDNA is oriented head-on with respect to fork progression (Figure 5B). Indeed, assays dependent on downstream DNA damage, as well as 2-Dimensional gels, may detect stalls or arrests that are below the limit of detection in our assay; however, our assay allows for the quantification and direct comparison of different classes of potential replisome impediment in the same samples. Therefore, while we cannot conclude from our analysis that G-quadruplexes and RNAP2 genes do not induce any replisome stalling or arrest in *pif1* and/or *rrm3* mutant cells, we establish that any such changes in fork mobility are extremely rare compared to those observed at tDNAs. We suggest that G-quadruplexes and/or highly transcribed RNAP2 genes may indeed cause DNA damage as previously described, but without necessarily impeding the mobility of the replication fork.

Some previous reports of tDNA-mediated replisome arrest suggested positive supercoiling ahead of the tDNA as a contributing mechanism due to relatively de-localised stalling by 2-D gel and a strong orientation bias^38^. Our high-resolution data indicate that both co-oriented and head-on collisions lead to significant fork arrest centered precisely on the tDNA (Fig. 4B). Although R-loops do not contribute significantly to the observed arrest (Fig. 6), the most parsimonious interpretation of our data is that the RNAP3 transcription complex itself represents an asymmetric barrier to the passage of the replication fork. We note that the leading edge of RNA polymerase has previously been identified as a replication barrier in prokaryotic systems^45^. Moreover, the observation that both Pif1 and Rrm3 suppress replisome arrest equivalently for both tDNA orientations lends further support to the characterization of these proteins as replisome-accessory ‘sweepases’. The inability of Rrm3 to remove a prokaryotic Tus-Ter replication block *in vivo*^46^ remains an unexplained exception to this model.

### Redundant Pif1 helicase activities and S-phase transcription

Fast-growing eukaryotes inevitably face conflicts between replication and transcription because the complete cessation of RNAP1 and RNAP3 transcription during S-phase – which accounts for one third of the 90-minute cell cycle in *S. cerevisiae*^47^ – would limit growth rate by reducing protein-synthesis capacity. Indeed, the ongoing high rate of tRNA synthesis during S-phase is exemplified by the immediate re-establishment of RNAP3 transcription complexes on nascent Okazaki fragments^18^. The *S. cerevisiae* rDNA repeats, transcribed by RNAP1, are unidirectionally replicated to avoid head-on collisions between replication and transcription^48^,^49^. Unlike the rDNA, tDNAs are dispersed throughout the *S. cerevisiae* genome and cannot use a directional RFB. Here we present the first evidence, to our knowledge, for the preferential orientation of eukaryotic tDNAs with respect to replication fork movement (Fig. 5). As in the rDNA, head-on collisions have been disfavored by evolution – presumably because they have a greater capacity to arrest replisomes than do co-directional collisions; alternatively, certain classes of replisome stalling or arrest might be more prone to cause DNA damage (Fig. 5B). However, we note that *S. cerevisiae* has maintained a significant fraction of tDNAs in the head-on orientation as opposed to eliminating head-on collisions entirely (Fig. 5E). It was recently proposed that *Bacillus subtilis* maintains a subset of essential genes in the normally disfavored head-on orientation in order to speed their evolution by stimulating replication-transcription conflicts^50^, which lead to TLS-dependent mutagenesis^51^. A plausible extension of this hypothesis to budding yeast would be that fork arrest at tDNAs could stimulate non-allelic homologous recombination between adjacent retrotransposons, thereby stimulating genomic rearrangements.

The lack of a requirement for RNAP3 transcription during S-phase in slower-growing cells of higher eukaryotes may underlie the extremely mild phenotypes observed in *PIF1* knockout mice^52^. However, metazoan Pif1 may have a more significant impact on replication dynamics and genome integrity in the context of transformed cells^53^. It has also been shown that RNA Pol II transcription is spatially separated from sites of active replication during S-phase^54^ but transcription by RNA Pol I and III was not evaluated. We also note that in an organism in which RNAP1 and RNAP3 transcription need not be maintained through S-phase, the only inevitable replication-transcription conflicts occur at extremely long genes (where transcription would take longer than a single cell cycle). Such long genes have indeed been identified as fragile sites in human cells^37^ but are not found in compact genomes such as *S. cerevisiae*. Fragility at these sites appears to R-loop dependent, but replisome stalling at these loci has not been directly investigated.

### Reconciling the absence of replication fork stalling with increased rates of genomic instability at G-quadruplexes and R-loops

Our data provide evidence against substantial, prolonged replisome stalling or arrest at G-quadruplexes, even in a *pif1-m2 rrm3Δ* strain grown in the presence of low concentrations of hydroxyurea (Fig 4 and S6D). However, evolutionarily diverged Pif1 helicases from both eukaryotes and prokaryotes preferentially unwind G-quadruplexes *in vitro*^10^. Moreover, mutation rate, microsatellite instability, and gross chromosomal rearrangements around model G-quadruplexes increase in the absence of Pif1 activity^10^,^28^,^32^. Therefore, G-quadruplexes are unstable in the absence of Pif1 even under conditions where they do not broadly impact replisome mobility. It is possible that fork blockage at G-quadruxplexes remains below the level of detection of our assay, but that techniques that select for rare events (such as a GCR screen) may identify biologically relevant but extremely rare fork impediments, or damage that is independent of replisome stalling. While predicted G-quadruplex sequences exist in the *S. cerevisiae* genome, it is unclear how many actually form stable intramolecular structures *in* vivo in the absence of G4-stabilizing drugs^55^. Even so, since Pif1 recruitment to G-quadruplexes has been reported to occur long after replisome passage^28^, we propose that genome instability due to these structures is separable from replication fork stalling, and may reflect a post-replicative repair phenomenon.

Similarly to G-quadruplexes, R-loops are associated with DNA damage, but our data suggest that they do not significantly stall the replication fork even in the absence of Rrm3 (Fig. 6). A genome-wide analysis of DNA damage hotspots by γH2A ChIP-chip found little sign of replication-stalling induced damage at sites that had been previously predicted to stall the fork^56^. Furthermore, recent experiments in human cells have demonstrated that R-loop structures are repaired via a NER-like pathway prone to double-stranded breaks that, importantly, do not require the collision of a replication fork and instead are the direct result of R-loop processing events^57^. Our data support a model in which replisome stalling and DNA damage due to both R-loop and G-quadruplex formation are mechanistically separable, and underscore the importance of using a damage-independent and genome wide assay, like Okazaki fragment sequencing, to evaluate replisome mobility, pausing, and arrest.

## EXPERIMENTAL PROCEDURES

### *In vivo* methods

All *S. cerevisiae* strains are derived from W303 *RAD5*+, and were grown in YEPD at 30°C unless otherwise noted. To minimize the accumulation of suppressors in *pif1-m2 rrm3Δ*, which has been reported to show elevated genomic instability^10^, we froze stocks of all strains after the minimum feasible number of divisions and conducted all experiments using freshly streaked cells from these stocks. The three wild-type, *rrm3Δ*, *pif1-m2* and *pif1-m2 rrm3Δ* strains used were biological replicates taken from segregants from three single tetrads following sporulation of a *pif1-m2/PIF1 rrm3/RRM3 CDC9*^*REP*^*/CDC9*^*REP*^ diploid. Strains that used a Gal1,10 promoter to drive the expression of *RNH1* were grown overnight in Galactose (2%) prior to ligase repression.

Following growth to mid-log phase, DNA ligase expression was repressed in asynchronous cultures by treatment with 40 mg/L doxycycline for 2.5h. For replication stress experiments, hydroxyurea was added to a final concentration of 25 mM 1h prior to addition of doxycycline, which was added to cultures at the same concentration for 2h. Okazaki fragments were labeled, purified, and deep-sequenced as previously described^18^. Paired end sequencing (2 x 50bp) was carried out on the Illumina Hi-seq 2500 platform.

### Computational methods

Reads were mapped to the S288c reference genome^58^ using Bowtie (v2.2.3) and converted into bamfiles using samtools, removing PCR duplicates with the MarkDuplicates function in picard tools. In-house python and command line scripts were used to convert to 1-based bedpe and sorted bedfiles, with the strand and 5’- and 3’-end of each read indicated. We used bedtools to convert bedfiles into stranded sgr files giving per base read coverage over the genome; Python was used to calculate the percent of total reads mapping to either the Watson or Crick strand. All calculations were carried out using un-smoothed data. 1 was added to all values to avoid dividing by 0. Datasets were internally normalized by calculating the percent reads mapping to the Watson or Crick strand (i.e. a value independent of read depth) and only combined after analysis when we calculated the grand mean and p-value for fork arrest (see below).

For the analysis of fork progression at sites of interest (tDNAs, RNA Pol II genes, G-quadruplex forming sequences, random genomic positions), we determined the average fork movement at that site (leftward or rightward) by the percent Watson strand hits within 1kb of the site (>50% indicates leftward-moving; <50% indicates rightward-moving). Random sites were generated by sampling lines from an sgr file to ensure equal coverage per base per chromosome. We calculated the percent of rightward-moving forks by the percent of reads mapping to the Crick strand. We then analyzed leftward- and rightward-moving forks separately so that we could reverse the direction of leftward-moving forks and sum the average percent of forks moving unidirectionally over a 10kB window of analysis (i.e. value = average(rev(percentWatson(left-to-right)) + (100-(percentWatson(right-to-left))), see Figure 2C and D and Figure S4A).

To calculate the percent change in fork direction at each site, we used a window 1-3kB upstream and downstream of the site (see Figure 2D). The change in mean reads mapping to the Watson strand over this window indicates the percent of forks that have stalled or arrested at this site (see Figure 2B). Bar graphs indicate the grand mean of three biological replicates and the standard deviation of the three independent means. For all classes of sites (i.e. random sites versus G-quadruplex sites), we excluded sites that have an origin or tDNA within the 10kB window of analysis or that have a 100bp bin with no data unless otherwise noted. Sites were binned into co-directional and head-on by the average fork direction within 1kb of the site, as described above, and the direction of transcription (or the strand of the G-quadruplex forming structure). For RNA Pol II genes, we took the top 10% of genes expressed in YEP^29^ as ‘highly transcribed,’ but had similar results with more or less stringent subsets. We calculated the termination signal at the transcription start site, midpoint, and transcription termination site of the gene, but only report the transcription start site as the data were similar (i.e. no significant differences between strains).

Drip-seq data^19^ were accessed via Geo and analyzed using an in-house pipeline. DRiP-seq signal around tDNAs was calculated using a 400bp window and tDNAs were sorted by their integrated DRiP-seq signal in WT (see Figure 6A).

### Statistical methods

We evaluated the significance of the change in replication direction between datasets and sites by Monte Carlo resampling. Briefly, we compared the grand mean of two treatments (e.g. WT versus *pif1-m2 rrm3Δ* at 93 tDNAs) before creating a dataset that conformed to the null hypothesis by randomly assigning data into the two treatments. Data were randomly assigned 10,000 times, and the difference in grand mean calculated for each sampling; the number of times that the resampled data (i.e. the null hypothesis) recreated the difference in means calculated for the data denoted the p-value. Monte Carlo resampling was done many times (at 10,000 trials per run) until a p-value was converged upon. For datasets with more than 93 sites, we randomly sampled 93 of the total sites before randomly assigning data into the two treatments; when the p-value varied widely between runs (due to different lines in the file being tested), data were considered insignificant (p>0.05) if more than half of the trials returned insignificant p-values; in these cases, we looked for smaller numbers of sites that were driving the potentially significant differences (see Figure S6E). In Figures, (*) denotes a significant p-value between 0.05 and 0.0002; (***) denotes a p-value less than 0.0001, wherein our many trials of 10,000 resamplings never produced a single run where the observed difference in means could be explained by random sampling.

### 2-Dimensional gel analysis

The plasmid for our 2-Dimensional gel analyses (Figure 3B) was cloned by standard techniques from pRS426, which contained a Ura marker and 2μ yeast origin of replication. We replaced the entire 2μ origin sequence with ARS309 and inserted either *Pgk1* or *tG(TCC)C* with 250bp of their native upstream promoter sequences to drive expression. 100ml cells were grown in -URA media to mid-log phase, collected by centrifugation, and resuspended in two 0.75ml aliquots in 50mM Tris HCl pH 8.0, 50mM EDTA, 100μg/ml psoralen before 12min crosslinking at 365nm. Cells were then pelleted, washed in 0.5ml TE, pelleted again and stored at −20°C until analysis.

DNA was purified by standard techniques. Briefly, cell pellets were resuspended in 0.5ml TE buffer with 1:100 β-ME and 250μg/ml zymolyase (T100) and incubated for 30 minutes before the addition of 100μl lysis buffer (500mM Tris HCl pH 8.0, 0.25M EDTA, 3% SDS) and 20μl Proteinase K (Roche). After 2h at 50°C, 150uL 5M potassium acetate was added and cell debris was pelleted for 30min at top speed at 4°C. The supernatant was precipitated with ethanol, washed, dried and resuspended in 750μl TE with RNase A (50μg/ml) for at least 1h at 37°C before phenol:cholorform extraction and isopropanol precipitation. Pellets were resuspended in 150μl TE o/n and digested o/n with EcoRI-HF (*tG(TCC)C* or empty vector) or EcoRV (*Pgk1*) in 600μl total volume with 400 units of enzyme. DNA was precipitated prior to loading on a 0.4% agarose gel run at room temperature (22V, 18-24h). Slices containing the size of interest were cut, turned counter-clockwise 90°, and run on a 0.95% agarose gel containing 0.3μg/ml Ethidium Bromide (130V, 18h, 4°C). Arcs were visualized, cut, nicked by autocrosslinking (Strata-linker), depurinated by treatment with acid (0.25N HCl), treated with denaturing solution (0.5N NaOH) and equilibrated in blotting solution (1.5MNaCl, 0.25N NaOH) before transfer onto Hybond N+ nylon membrane o/n. Blots were neutralized in 50mM sodium phosphate (pH 7.2), equilibrated at 65°C in hybridization buffer (0.25M Na phosphate pH 7.2, 0.25 M NaCl, 1mM EDTA, 7% SDS, 5% Dextran Suflate) before the addition of radiolabeled (Invitrogen Random Primers DNA Labeling Kit) probe (against the Amp^r^ gene of the plasmid, see Figure 3B). Blots were washed 5× 100ml in low (2X SSC, 0.1%SDS) and high (0.1X SSC, 0.1% SDS) stringency washes, patted dry, and exposed to phosphoimager screens. 2-demensional gel analyses are representative of at least two independent experiments.

## ACCESSION NUMBERS

Raw and processed sequencing data have been deposited to the Gene Expression Omnibus under accession number GSE71973.

## AUTHOR CONTRIBUTIONS

J.S.O., J.K. and R.Y. generated data; J.S.O., J.K. and D.J.S. analyzed data and interpreted results; J.S.O. and D.J.S. wrote the manuscript with input from J.K. and R.Y.

## ACKNOWLEDGEMENTS

We thank Ginger Zakian and members of the Zakian lab for the 2-D gel protocol, for communicating data prior to publication, and for helpful discussions. We additionally thank Sevinc Ercan, Andreas Hochwagen, Hannah Klein, and members of the Smith lab for insightful discussions and critical reading of the manuscript, Dan Tranchina for help with statistical analyses, and Viji Subramanian for assistance with 2-D gel electrophoresis. This work was supported by NIH grant R01 GM114340, a March of Dimes Basil O’Connor Starter Scholar award (FY15-BOC-2141) and the Searle Scholars program (all to D.J.S). J.S.O is supported by an American Cancer Society - New York Cancer Research Fund postdoctoral fellowship (PF-16-096-01-DMB).

---

**Figure S1:**
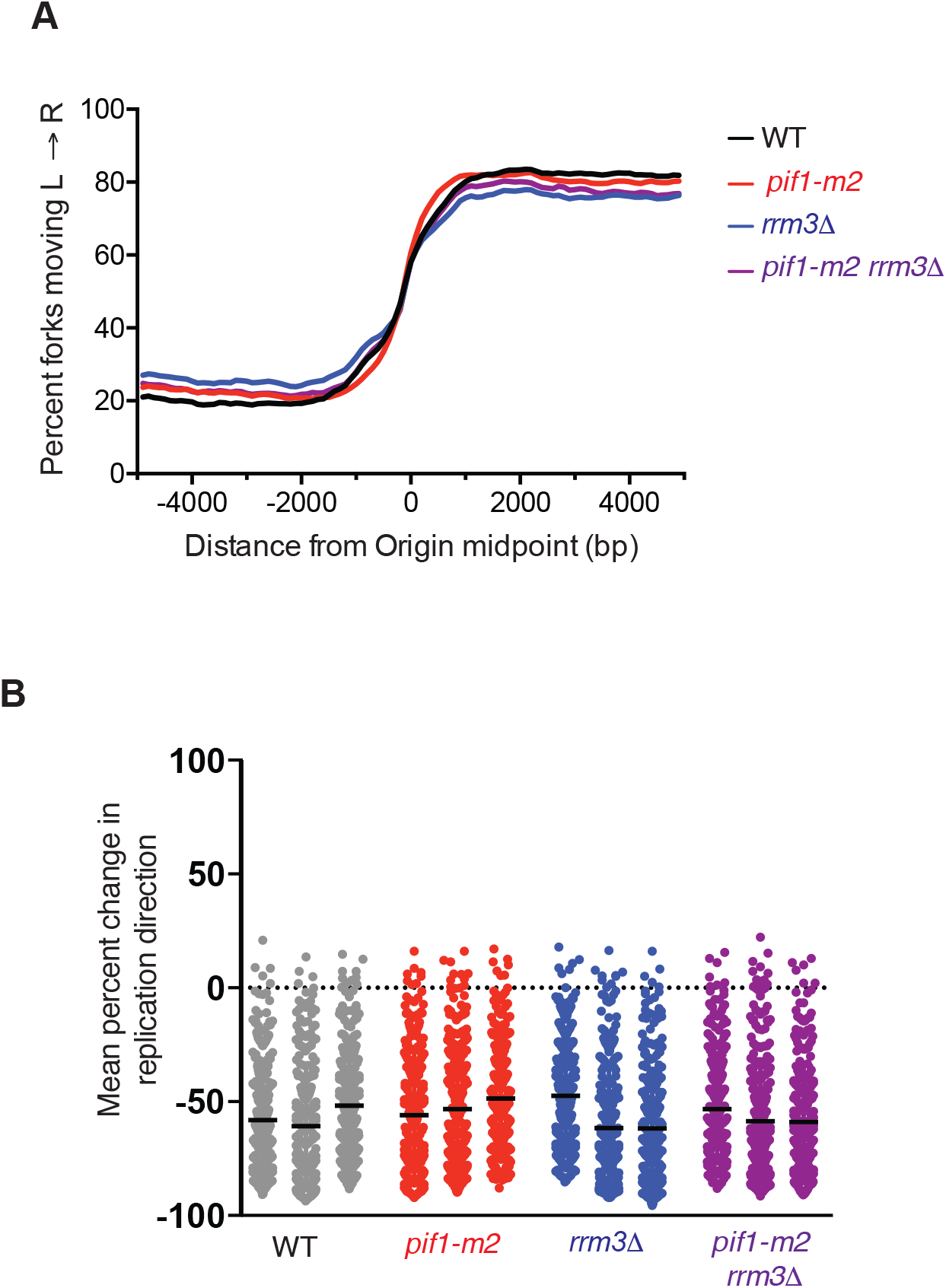
Origin use and efficiency is similar between WT, *rrm3Δ*, *pif1-m2*, and *pif1-m2 rrm3Δ* strains. A. Analysis of fork progression around confirmed and likely origins (McGuffee *et al.,* 2013). The percent of forks moving from left to right was calculated by the proportion of reads mapping to the Crick strand for each 100bp bin around the annotated origin of replication averaged across all sites (n=265). The origin signal is an ascending slope in this graph (compare to Figure 2A(i) and C). B. Scatter plot of origin usage at all confirmed and likely origins (McGuffee *et al.,* 2013). The origin signal was calculated at each site by the difference between the percent of forks moving right to left upstream (+1-3kb) and downstream (-1-3kb) of the site, similar to the termination signal described in Figure 2, but with the opposite polarity.

**Figure S2:**
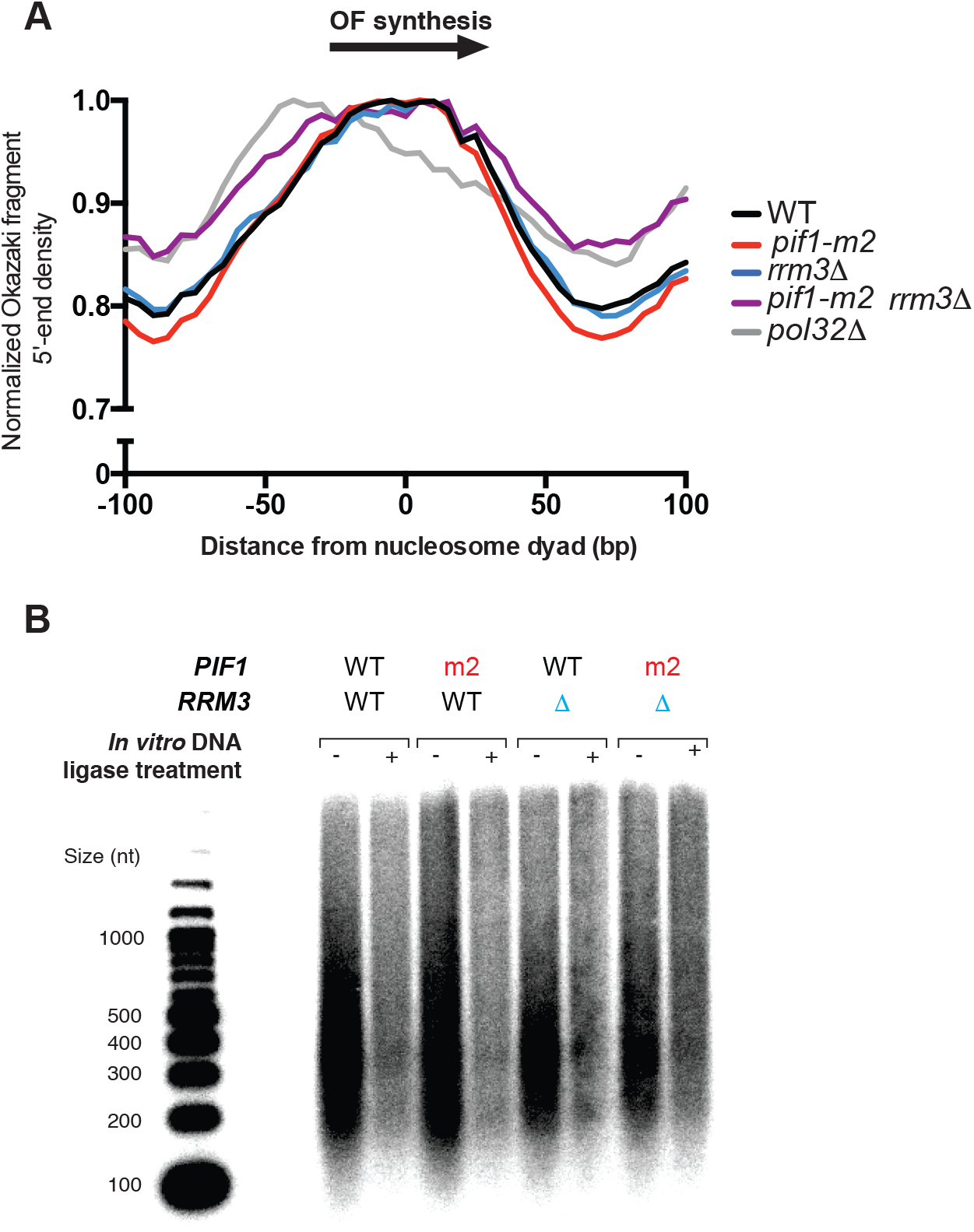
Further analysis of Okazaki fragment termini in pif1 and rrm3 mutant strains. A. Distribution of Okazaki fragment 3’ termini around nucleosome dyads (Jiang and Pugh, 2009) as shown for 5’-ends in Fig 1B. B. *rrm3Δ*, *pif1-m2*, and *pif1-m2 rrm3Δ S. cerevisiae* strains generate fully ligatable, nucleosome-sized Okazaki fragments similar to those observed in wild-type cells. Genomic DNA was prepared and labeled as described in the Materials and Methods.

**Figure S3:**
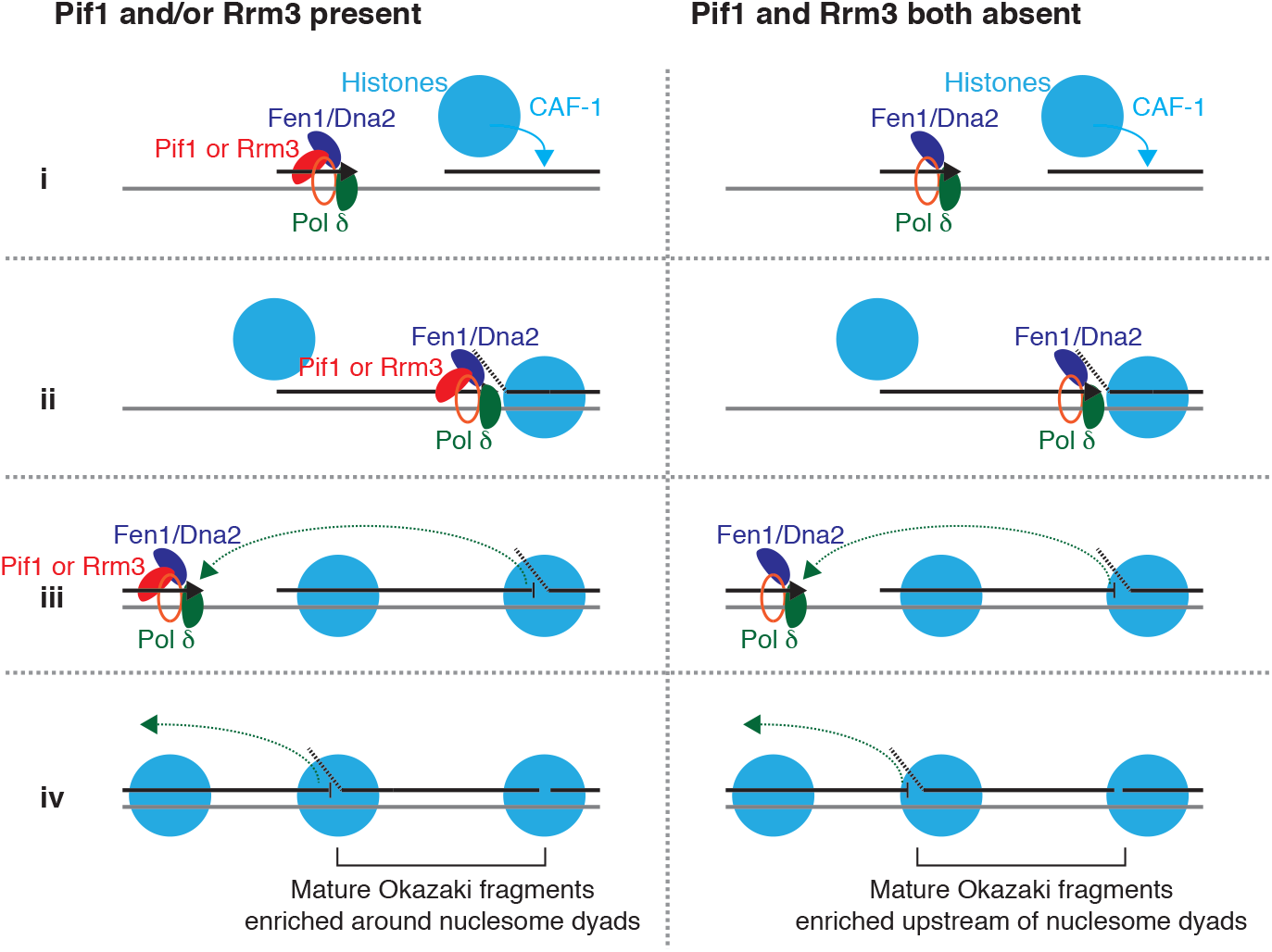
Model for Pif1 and Rrm3 activity on the lagging strand. Histones are redeposited on the nascent lagging strand through the action of Caf1 (blue) as the lagging strand is being. Either Pif1 or Rrm3 (red) is required for normal DNA Pol δ (green) processivity, with Okazaki fragment ends enriched around nucleosome dyads as shown in Figure 1B. In the absence of both Pif1 and Rrm3, mature Okazaki fragment ends are enriched upstream of the nucleosome dyads.

**Figure S4:**
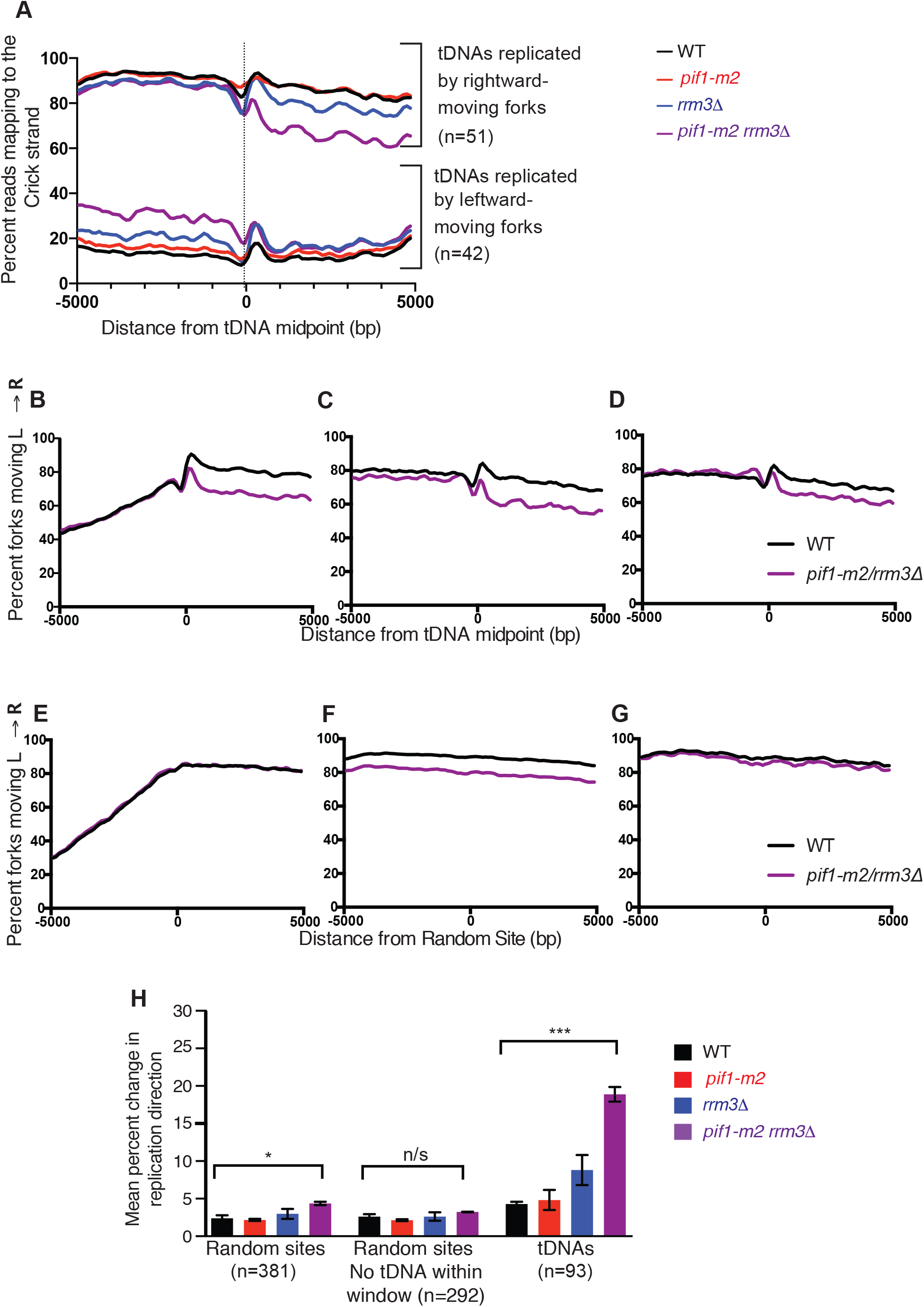
tDNAs are the predominant site of fork stalling genome wide. A. Fork progression of rightward (upper) and leftward (lower) moving forks at the 93 tDNAs selected for further analysis (see Materials and Methods). The percent of forks moving left-to-right was calculated by the percent of reads mapping to the Crick strand. B. Fork progression at all tDNAs with an origin in the analysis window in the WT (black) and *pif1-m2 rrm3Δ* (purple) mutant cells. The percent of forks moving left-to-right was calculated as described in the Materials and Methods with a correction for the slight difference in origin efficiency in the *pif1-m2 rrm3Δ* mutant (see Figure S1). The ascending slope in the upstream region is the origin signal (see Figure S1A). C. Fork progression at all tDNAs without an origin within the analysis window as shown in part B. D. Fork progression at tDNAs without an origin in the analysis window and without and origin-proximal tDNA plotted as in part B, and corrected for the slight difference in origin efficiency. E-G. Random sites with an origin within the analysis window (E), without an origin in the analysis window (F), and without an tDNA in the analysis window or proximal to the nearest upstream origin of replication (G) plotted as in part B with a correction for the slight difference in origin efficiency. H. Grand mean ± SD of the change in replication direction, interpreted as indicative of replisome stalling, at random sites, random sites that exclude tDNAs in the window of analysis, and tDNAs in three independent biological replicates for the indicated strains. Significance was determined by Monte Carlo resampling; *** indicates a p<0.0001; n/s indicates p>0.05.

**Figure S5:**
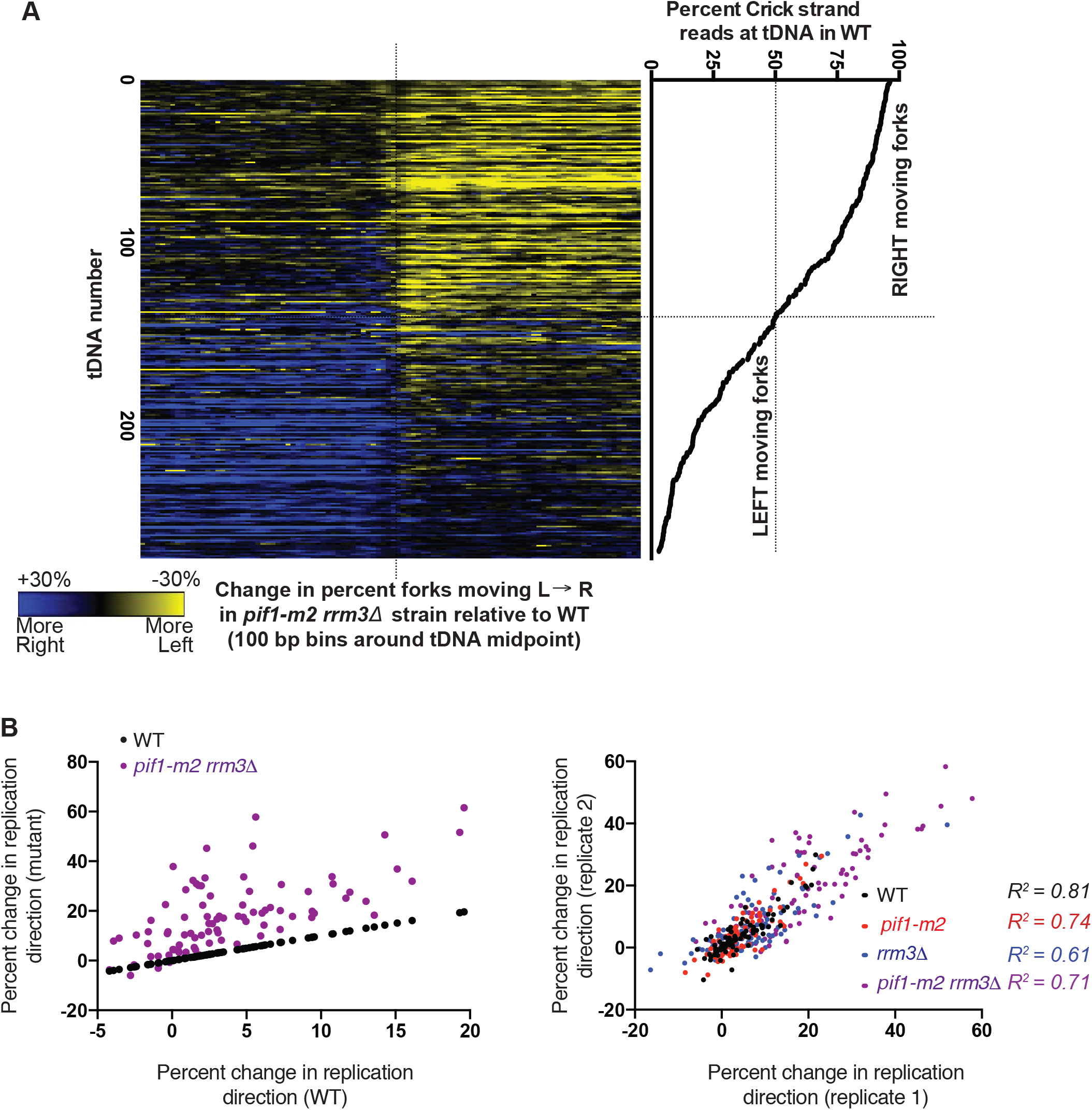
Fork arrest is a feature common to tDNAs. A. Heatmap showing the difference in percent forks moving left to right (calculated by the percent reads mapping to the Crick strand) between the WT and the *pif1-m2 rrm3Δ* double mutant. tDNAs were sorted by their average fork direction (see Materials and Methods), and data were binned to 100bp. Negative values (i.e. *pif1-m2 rrm3Δ* forks more leftward moving than wild type) were visualized in yellow while positive values (i.e. *pif1-m2 rrm3Δ* forks more rightward moving than wild the) were visualized in blue. Heatmap was constructed with Gitools. B. Scatter plot of all 93 tDNAs without a nearby origin or sequence gap. The change in replication direction was calculated for the same site in the WT and *pif1-m2 rrm3Δ* strains. Black dots are the WT plotted against the WT data and therefore give a line with the slope of 1. The *pif1-m2 rrm3Δ* change in replication direction divided by the change in the WT strain is plotted in purple. C. Reproducibility of tDNA arrest signal from biological replicate 1 to biological replicate 2. The change in replication direction from dataset 1 and dataset 2 at the 93 tDNAs included in our analysis. Each site was plotted four times, once for each strain, and the data were colored by strain. R^2^-values for the correlation between datasets for each strain were calculated using Graphpad Prism.

**Figure S6:**
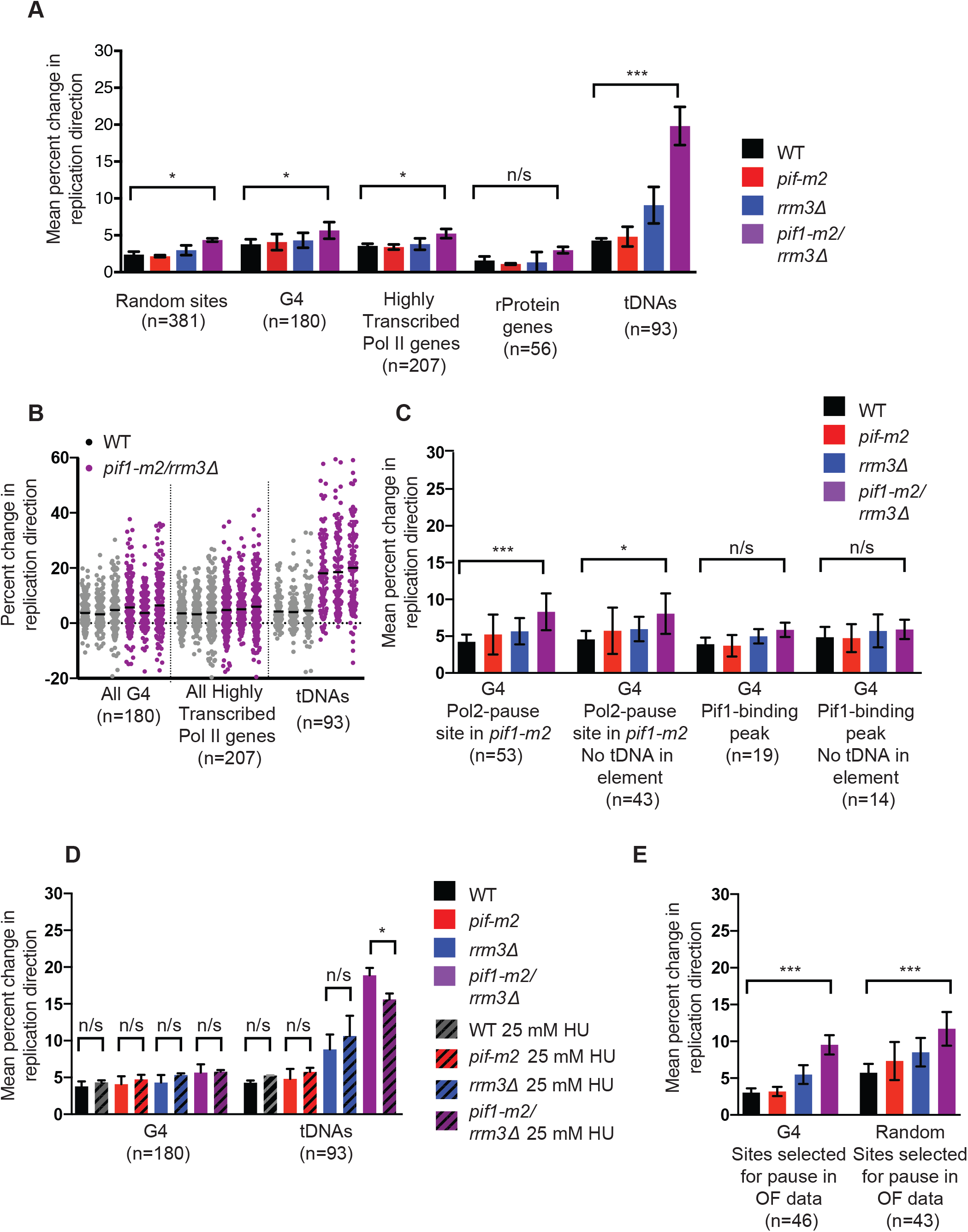
Lack of robust stalling or arrest signal at RNA Pol II genes and G-quadruplex forming sequences. A. Grand mean ± SD of the change in replication direction, interpreted as indicative of replisome stalling or arrest, at G-quadruplex forming sequences (Capra *et al.,* 2010), Highly transcribed RNA Pol II genes (Pelechano *et al*, 2013), ribosomal protein genes, and tDNAs as calculated in Figure 2E. In this analysis, random sites, G-quadruplex sequences, and RNA pol II genes with a tDNA in the analysis window were not removed (compare to Figure 4A). Significance was determined by Monte Carlo resampling; *** indicates a p<0.0001; * indicates 0.0001 < p < 0.05. n/s indicates p>0.05. B. Scatter plot of the change in replication direction at individual sites for all G-quadruplex forming sequences, highly transcribed RNA pol II genes, and tDNAs for the WT (gray) and *pif1-m2 rrm3Δ* strain (purple). Sites are as defined in part A. Individual termination signals were calculated at each site as described in Figure 2 and the sites used were described in part A. The mean of each dataset is depicted as a black bar. C. Grand mean ± SD of the change in replication direction from three biological replicates at different subsets of G-quadruplex sites previously shown to stall the fork or bind to Pif1 (Paeschke). Significance was determined by Monte Carlo resampling; *** indicates a p<0.0001; * indicates 0.0001 < p < 0.05. n/s indicates p>0.05. D. Selection of a subset of G-quadruplexes and random sites with a significant stalling in the *pif1-m2 rrm3Δ* double mutant. For both the G-quadruplex forming sequences (p<0.0001) and the random sites (p<0.0001), we were able to select a subset of the sites that shows significant stalling. Significance was determined by Monte Carlo resampling; *** indicates a p<0.0001; * indicates 0.0001 < p < 0.05. n/s indicates p>0.05. E. Addition of Hydroxy Urea (HU, 25mM) to the growth media, which slows replication forks by depleting the dNTP pool, did not increase specific stalling at G-quadruplex forming structures or tDNA sites. Grand mean ± SD of the change in replication direction, interpreted as indicative of replisome stalling, was plotted for G-quadruplex forming sites and tDNAs as in part A. Significance was determined by Monte Carlo resampling; *** indicates a p<0.0001. n/s indicates p>0.05.

**Figure S7:**
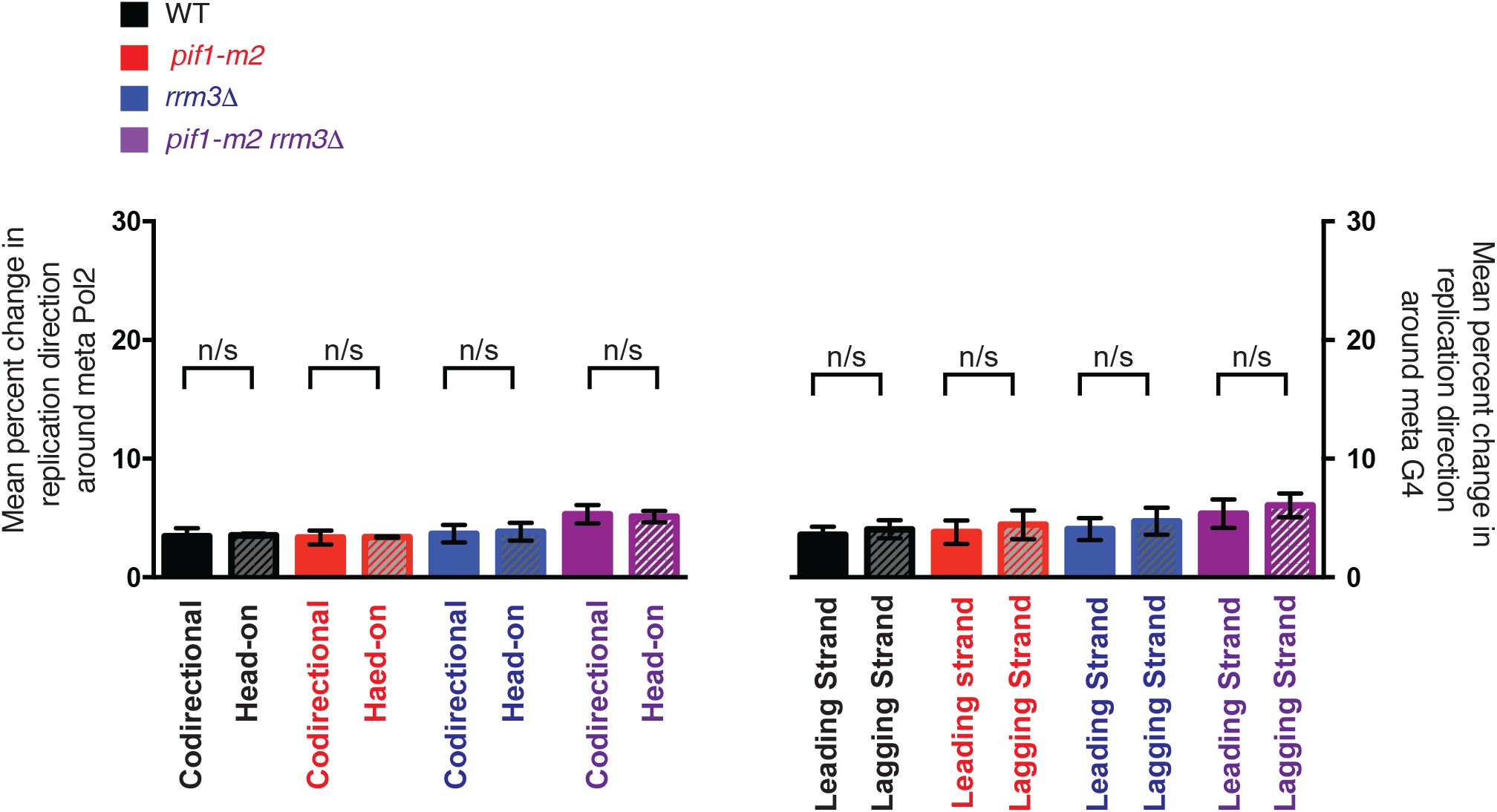
Direction of transcription at RNA Pol II genes and the strand of the G-quadruplex structure do not affect fork stalling or arrest at these sites. A. Grand mean ± SD of the change in replication direction, interpreted as indicative of replisome stalling or arrest, at RNA Pol II genes and G-quadruplex sequences binned by direction of replication (see Materials and Methods). Significance was determined by Monte Carlo resampling; n/s indicates p>0.05.

